# Spatial proteomic profiling of cardiac stromal populations and microenvironments in systemic sclerosis-associated primary myocardial involvement

**DOI:** 10.64898/2026.10.17.758293

**Authors:** Ayla Nadja Stütz, Giacomo de Luca, Alexandru-Emil Matei, Yi-Nan Li, Tim Filla, Veronica Batani, Bilgesu Safak Tümerdem, Aleix Rius Rigau, Cosimo Bruni, Wolfgang Merkt, Elric Zweck, Maike Büttner-Herold, Stefania Rizzo, Giovanni Peretto, Monica De Gaspari, Markus Eckstein, Georg Schett, Cristina Basso, Marco Matucci-Cerinic, Jörg H. W. Distler, Andrea-Hermina Györfi

**Author notes:** These authors contributed equally (Marco Matucci-Cerinic, Jörg H. W. Distler, Andrea-Hermina Györfi). These authors contributed equally (Ayla Nadja Stütz, Giacomo de Luca). **Corresponding authors:** Jörg H.W. Distler, MD, Department of Rheumatology and Hiller Research Center, University Hospital Düsseldorf, Heinrich-Heine University, 40225 Düsseldorf, Germany, Andrea-Hermina Györfi, MD, Department of Rheumatology and Hiller Research Center, University Hospital Düsseldorf, Heinrich-Heine University, 40225 Düsseldorf, Germany.

## Abstract

**Objectives:** To define cardiac stromal heterogeneity and spatial stromal-immune organization in systemic sclerosis-associated virus-negative myocarditis (SSc-VNM), a manifestation of primary SSc heart involvement.

**Methods:** Imaging mass cytometry, a single-cell multiplexed spatial proteomic technology, was applied to endomyocardial biopsies from ten patients with SSc-VNM, seven with VNM associated with other systemic autoimmune rheumatic diseases (a-VNM) and nine with isolated VNM (i-VNM). Single-cell phenotyping, cellular-neighborhood and spatial-interaction analyses, and correlations with histological fibrosis were performed at patient level.

**Results:** Among 82,673 segmented cells, we identified seven fibroblast phenotypes and one CD34^+^ stromal-cell population. The overall fibroblast/stromal compartment was expanded in SSc-VNM compared with i-VNM (adjusted p = 0.00973). The composition of fibroblasts/stromal cells differed profoundly in SSc-VNM compared with a-VNM and i-VNM, with numerical increases in CD34^+^ stromal cells and SOX9^+^ and PI16^high^/POSTN^high^ fibroblasts and de-enrichment of FAP^high^/POSTN^high^ fibroblasts and myofibroblasts. Spatial analysis identified ten cellular neighborhoods and revealed distinct SSc-VNM-associated patterns, including altered stromal-immune interaction networks. These included enriched interactions of CD34^+^ stromal cells and PI16^high^/POSTN^high^ fibroblasts with CD163^high^/HLA-DR^+^ macrophages and CD68^+^ mast cells. PI16^high^/POSTN^high^ and PI16^-^/POSTN^high^ fibroblasts demonstrated the highest collagen type 1 expression among all stromal cell subsets across all VNM entities and PI16^high^/POSTN^high^ fibroblasts were numerically increased in biopsies with ≥10% fibrotic remodeling and were spatially associated with collagen-rich areas.

**Conclusions:** Our findings refine current concepts of human fibrotic myocardial remodeling and support therapeutic targeting of specific stromal players such as POSTN^high^ fibroblast subpopulations.

**Key messages:** *What is already known on this topic:*

- Primary SSc heart involvement contributes substantially to disease-related mortality, but its pathophysiology remains largely enigmatic, particularly lacking understanding of cellular interactions and spatial architecture.
- Single-cell studies have established fibroblast heterogeneity in both human cardiac disease and SSc skin and lung tissues.

*What this study adds:*

- We provide a molecular characterization of cardiac stromal cell subpopulations using protein-based definitions and demonstrate dysregulation of several fibroblast/stromal subsets, with increases in CD34^+^ stromal cells and SOX9^+^ and PI16^high^/POSTN^high^ fibroblasts and de-enrichment of FAP^high^/POSTN^high^ fibroblasts in SSc-VNM compared with a-VNM and i-VNM.
- We identify the SOX9^+^ fibroblast niche as a distinct microenvironment particularly enriched in immune cells, including monocyte and macrophage subsets.
- We demonstrate major differences in the spatial cellular interaction networks of cardiac fibroblasts/stromal cells in SSc-VNM compared with a-VNM and i-VNM.
- We identify PI16^high^/POSTN^high^ and PI16^-^/POSTN^high^ fibroblasts as the main collagen type 1 producing fibroblast subsets. Consistently, PI16^high^/POSTN^high^ fibroblasts were numerically increased in biopsies with ≥10% fibrotic remodeling and were associated with collagen-rich areas.

*How this study might affect research, practice or policy:*

- Our study identifies defined cardiac stromal states and stromal-immune niches as potential candidates for future therapeutic intervention in primary SSc heart involvement.

## Introduction

Fibrotic remodeling is a common response to myocardial injury and a major determinant of ventricular dysfunction and heart failure. Although cardiac fibroblasts are central effectors, they are not a uniform cell population. Human single-cell and spatial studies have resolved injury-associated fibroblast states with distinct matrix, inflammatory, vascular-support and progenitor-like programs across myocardial infarction and chronic cardiomyopathies.[1–5] These observations challenge a model, in which α-smooth-muscle-actin-positive myofibroblasts alone account for extracellular-matrix production and instead emphasize context-dependent stromal ecosystems. How these states are organized in autoimmune myocarditis remains incompletely defined.

Systemic sclerosis (SSc) is a systemic autoimmune connective tissue disease, which combines immune dysregulation, vasculopathy and progressive fibrosis.[6] Primary SSc heart involvement (SSc-pHI) is a major contributor to SSc-related mortality.[7–9] Fibrosis, inflammatory infiltrates and microvascular alterations frequently coexist in myocardial tissue, with SSc-associated virus-negative myocarditis (SSc-VNM) showing extensive matrix deposition and adverse outcomes.[10,11] Single-cell and spatial analyses have meanwhile resolved pathogenic fibroblast and immune niches in SSc skin and lung tissues.[12–14] Whether analogous cell states and spatial microenvironments shape the myocardium in SSc-pHI has not been examined at single-cell protein resolution.

Recent bulk transcriptomic profiling of SSc myocardial biopsies identified suppression of mitochondrial energy metabolism, enhanced autophagy and macrophage-associated inflammatory programs.[15] However, this technique does not provide single-cell or spatial resolution. We used imaging mass cytometry (IMC) to map cardiac fibroblast states, cellular neighborhoods and stromal-immune interactions in endomyocardial biopsies (EMBs) from patients with SSc-VNM. Virus-negative myocarditis associated with other systemic autoimmune rheumatic diseases (a-VNM) and isolated virus-negative myocarditis (i-VNM) provided disease controls. We further examined how spatial features related to histological myocardial fibrosis and would merit targeted validation in future studies.

## Methods

### Study design and participants

This multicenter study analyzed diagnostic right-ventricular septal endomyocardial biopsies obtained between 2011 and 2018 from ten patients with SSc-VNM, seven with a-VNM and nine with i-VNM (figure 1A; supplementary table S1). Myocarditis was centrally adjudicated using the Dallas criteria, and cardiotropic viruses were excluded by quantitative PCR.[16] SSc was classified using the 2013 ACR/EULAR criteria and SSc-pHI was diagnosed according to the World Scleroderma Foundation/Heart Failure Association consensus definition.[8,17] All participants provided written informed consent and the study was approved by the ethical committees of the Medical Faculty of the Heinrich-Heine University in Düsseldorf (2022-2189), the Medical Faculty of the Vita-Salute San Raffaele University in Milan (22/INT/2018, PP_019/2), and the Medical Faculty of the University Hospital Erlangen in Erlangen (98_18 B, 30_19B).

**Figure 1.**
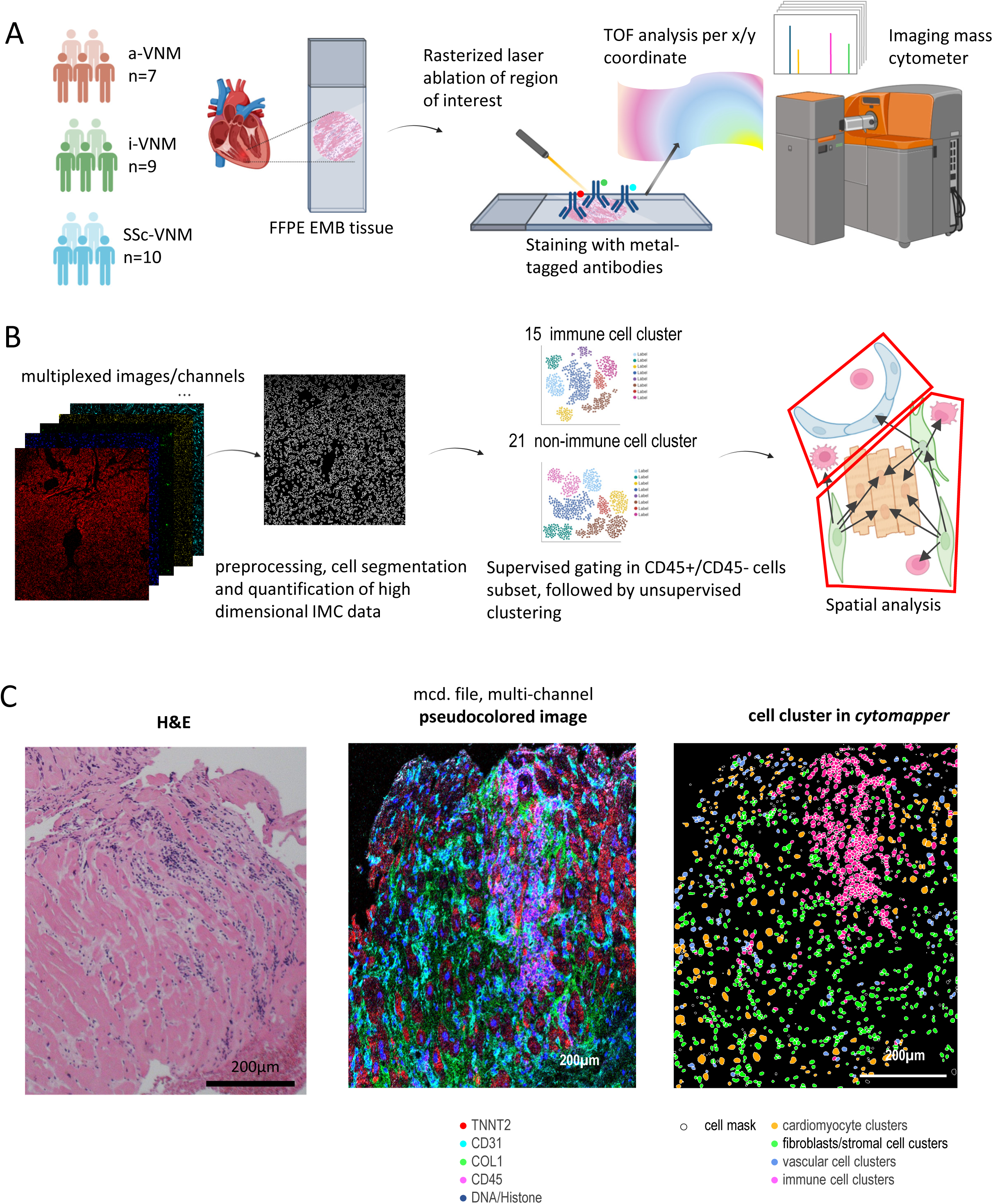
Imaging mass cytometry of endomyocardial biopsies in virus-negative myocarditis. (A) Cohorts and workflow: SSc-VNM (n=10), a-VNM (n=7) and i-VNM (n=9). Formalin-fixed, paraffin-embedded biopsies were stained with metal-tagged antibodies and acquired at 1 µm² pixel resolution. (B) Analysis workflow comprising image quality control, Steinbock preprocessing, Mesmer segmentation, supervised CD45 gating, PhenoGraph clustering, phenotypic annotation and spatial analyses. (C) From left to right: representative haematoxylin and eosin section; multiplex IMC image (TNNT2, red; CD31, cyan; COL1A1, green; CD45, magenta; DNA/histone, dark blue); and cytomapper image showing main cell clusters. a-VNM, virus-negative myocarditis associated with a non-SSc systemic autoimmune rheumatic disease; EMB, endomyocardial biopsy; i-VNM, isolated virus-negative myocarditis; SSc-VNM, systemic sclerosis-associated virus-negative myocarditis; TOF, time of flight. Some parts of the image were created with BioRender. Stütz, A. (2025) https://BioRender.com/pfaazx6, https://BioRender.com/5wakkhu

### Imaging mass cytometry and cell phenotyping

Five-micrometer formalin-fixed sections were stained with a validated metal-conjugated antibody panel (supplementary table S2). Regions of approximately 1.5 mm² were acquired at 1 µm² resolution, preprocessed with Steinbock and segmented with Mesmer (figure 1B).[18] After quality control and normalization, a random-forest classifier separated immune from non-immune cells; RPhenograph clustering and canonical marker combinations defined immune, vascular and stromal phenotypes.

### Spatial and statistical analyses

Four-nearest-neighbour graphs, 1000 label permutations and k-means clustering of local cell composition quantified pairwise interactions and cellular neighborhoods.[12] Patient was the inferential unit. Group differences were tested with Kruskal-Wallis/Dunn procedures or Propeller, with Benjamini-Hochberg correction as appropriate. For the primary group comparisons, samples were stratified by histological fibrosis burden, defined as low (<10% extracellular matrix) or moderate/severe (≥10% extracellular matrix). In the absence of unequivocally validated thresholds for myocardial fibrosis burden, these cut-offs were selected arbitrarily. Complementary Spearman analyses assessed associations between the continuous extent of fibrosis and patient-level features; these correlations were considered exploratory after correction for multiple testing. Detailed laboratory, image-processing, phenotyping and statistical procedures are provided in the supplementary methods.

## Results

### Cohort and IMC-based cardiac cell phenotyping

The cohort comprised 26 patients. Epidemiological, clinical and histopathological data of the patients are presented in supplementary table S1.

IMC-based phenotyping yielded 82,673 segmented cells: 35,706 from SSc-VNM, 20,551 from a-VNM and 26,416 from i-VNM and included 17,985 stromal cells, 19,398 vascular cells and 6,444 immune cells. Segmentation and phenotype maps preserved myocardial architecture and enabled separate clustering of CD45^-^ non-immune and CD45^+^ immune compartments (figures 1 and 2; supplementary figures S1-S4).

Seven fibroblast phenotypes were resolved. Three of these fibroblast populations expressed high levels of periostin (POSTN), but differed clearly in the expression of other markers, and were annotated as PI16^-^/POSTN^high^, PI16^high^/POSTN^high^, and FAP^high^/POSTN^high^ fibroblasts, respectively. Additional fibroblast clusters were defined by TCF21, THY1/ADAM12/FSP1 or SOX9 (TCF21^+^, THY1^+^/ADAM12^+^/FSP1^+^, and SOX9^+^ fibroblasts). An additional cluster of α-smooth-muscle-actin (α-SMA)^high^ fibroblasts was annotated as myofibroblasts. A distinct stromal population expressed lower levels of PI16 and PDGFRα and of broader fibroblast markers compared to the other fibroblast populations but was characterized by high CD34 expression and was annotated therefore as CD34^+^ stromal cells (figure 2 and supplementary figure S5). Patient-level marker distributions showed the strongest COL1 signal in PI16^high^/POSTN^high^ and PI16^-^/POSTN^high^ fibroblasts, followed by myofibroblasts, supporting functional heterogeneity within the matrix-producing compartment (figure 2 and supplementary figure S5).

**Figure 2.**
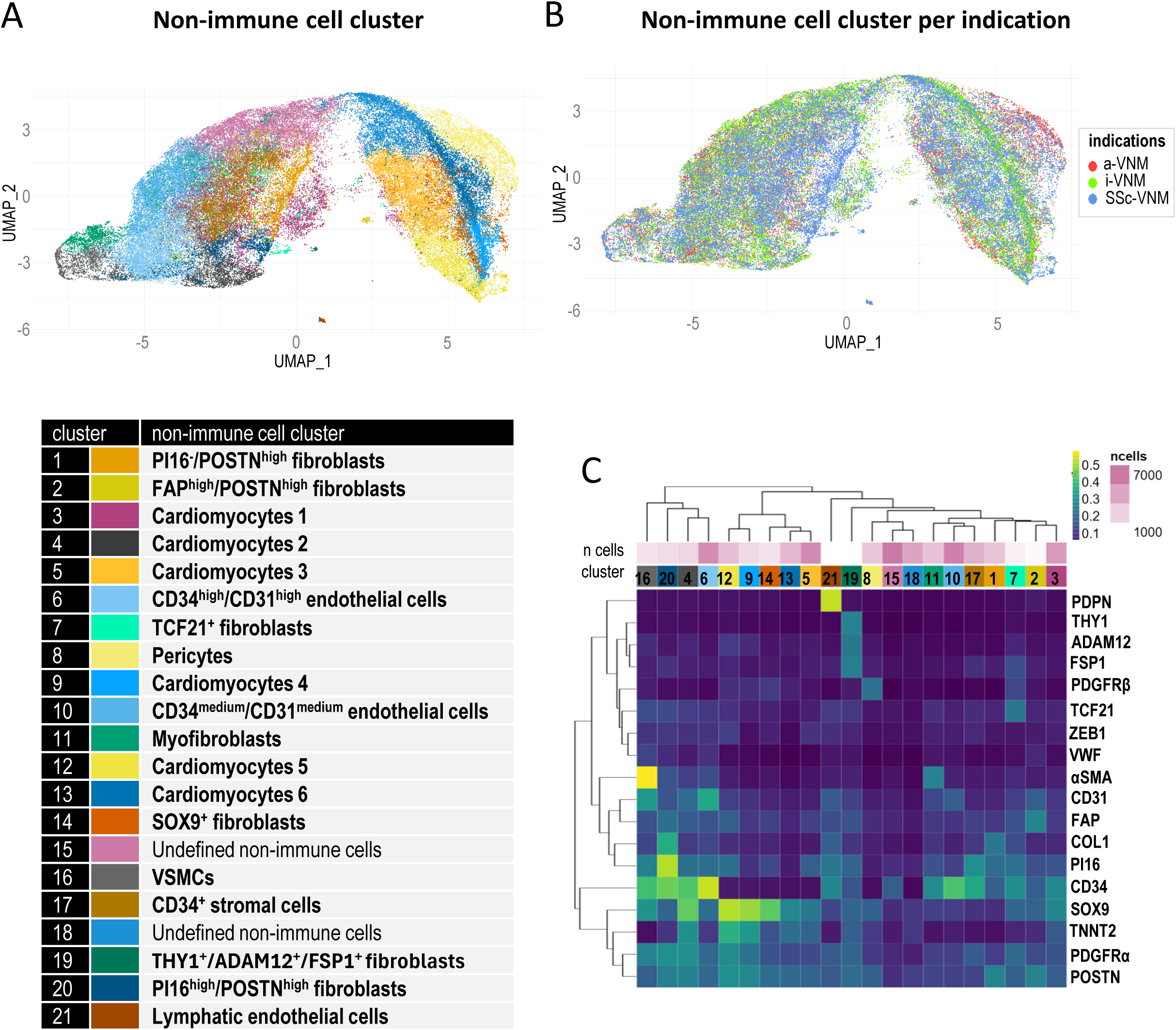
Phenotypic characterization of CD45^-^ non-immune populations. (A) Uniform Manifold Approximation and Projection for Dimension Reduction (UMAP) of clusters 1-21 coloured by annotated phenotype. (B) UMAP coloured by disease group. (C) Heatmap of scaled marker expression across clusters. EC, endothelial cell; LEC, lymphatic endothelial cell; UMAP, uniform manifold approximation and projection; VSMC, vascular smooth muscle cell; a-VNM, virus-negative myocarditis associated with a non-SSc systemic autoimmune rheumatic disease; i-VNM, isolated virus-negative myocarditis; SSc-VNM, systemic sclerosis-associated virus-negative myocarditis;

The vascular compartment comprised CD34^high^/CD31^high^ and CD34^medium^/CD31^medium^ endothelial cells (ECs), podoplanin (PDPN)^high^ lymphatic endothelial cells (LECs), PDGFRβ^+^ pericytes and α-SMA^high^ vascular smooth-muscle cells (VSMCs, figure 2 and supplementary figure S2).

Immune phenotyping resolved three macrophage states (CD163^high^/HLA-DR^+^ macrophages, CD163^+^/HLA-DR^-^ macrophages, HLA-DR^-^ macrophages), two monocyte states (CD86^high^ monocytes and HLA-DR^high^ monocytes), CD4^+^ T cells and CD8^+^ T cells, CD20^+^ B cells and HLA-DR^high^ lymphocytes (supplementary figure S3). A rare TPSAB1^+^/CD68^+^ cluster was retained after spatial review of separate and merged channels and annotated as CD68^+^ mast cells (supplementary figure S6), consistent with a protein-defined phenotype reported in colorectal-cancer stroma.[19]

### Disease-associated remodeling of stromal and immune compartments

Broad compartment composition differed across diseases (figure 3A, supplementary figure S7). Fibroblasts/stromal cells represented a mean 28.2% of cells in SSc-VNM, 21.6% in a-VNM and 13.6% in i-VNM. Patient-level fibroblast/stromal proportions were higher in SSc-VNM than in i-VNM (adjusted p (p_adj_) = 0.00973), while SSc-VNM versus a-VNM did not reach adjusted significance (figure 3A). Fibroblast/stromal-cell density was likewise higher in SSc-VNM than in i-VNM (p_adj_ = 0.015). Total cell density and the relative vascular and immune compartments were comparable, indicating selective fibroblast/stromal expansion rather than a uniform increase in tissue cellularity (supplementary figure S7).

**Figure 3.**
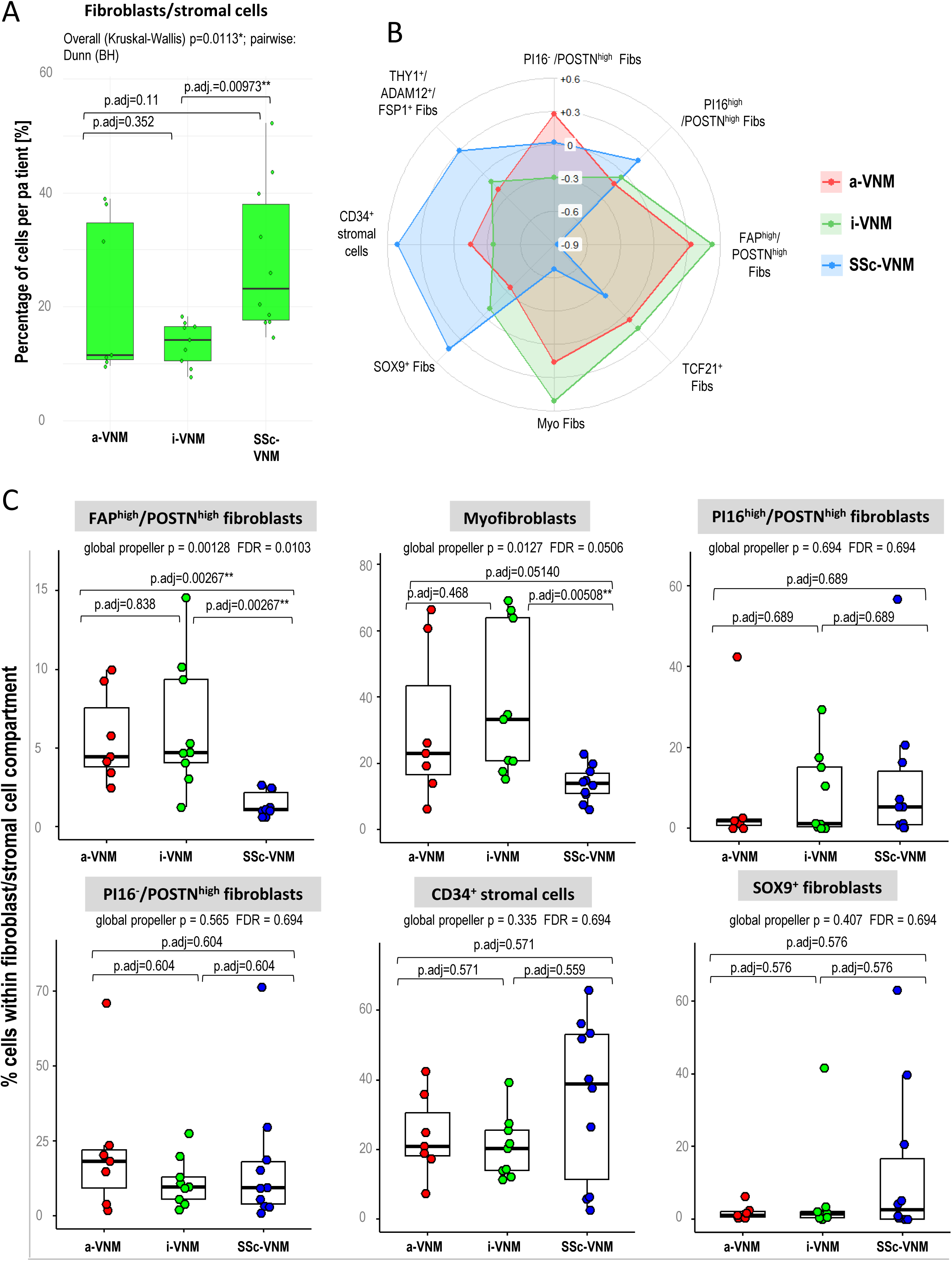
Disease-associated differences in cellular compartments and fibroblast/stromal populations. (A) Stromal-cell proportion per patient across myocarditis entities. Kruskal-Wallis p=0.0113; Dunn-Benjamini-Hochberg-adjusted pairwise p values are shown. (B) Disease-balanced z-scaled radar chart of stromal phenotypes; zero denotes the mean of disease-level means for each phenotype. (C) Patient-level proportions of selected phenotypes within the stromal compartment. Boxplots show medians, interquartile ranges and 1.5×IQR whiskers. Propeller global p values, FDR values and adjusted pairwise p values are displayed. a-VNM, virus-negative myocarditis associated with a non-SSc systemic autoimmune rheumatic disease; i-VNM, isolated virus-negative myocarditis; SSc-VNM, systemic sclerosis-associated virus-negative myocarditis.

The expanded SSc-VNM fibroblast/stromal compartment showed a distinctive balance of cellular populations (figure 3B-C and supplementary figure S8A). FAP^high^/POSTN^high^ fibroblasts differed across diseases (global FDR=0.0103) and were reduced in SSc-VNM (mean 1.43%) compared with a-VNM (5.66%) and i-VNM (6.35%; p_adj_ = 0.00267 for both). Myofibroblasts were also less frequent in SSc-VNM (13.88%) than in i-VNM (38.0%; p_adj_ = 0.00508). Together, these findings establish that greater fibroblast/stromal representation in SSc-VNM is not driven by uniform expansion of conventional activated fibroblast phenotypes when compared with other myocarditis disease entities (figure 3B-C).

Several additional populations formed coherent SSc-VNM-associated patterns. CD34^+^ stromal cells were the largest stromal population in SSc-VNM (mean 34.67% of stromal cells) and were numerically more abundant than in a-VNM (24.0%) and i-VNM (20.68%) (supplementary figure S8A). The number of SOX9^+^ fibroblasts was also increased in SSc-VNM (13.41% in SSc-VNM, 1.83% in a-VNM and 5.77% in i-VNM) (supplementary figure S8A). PI16^high^/POSTN^high^ fibroblasts were modestly enriched in SSc-VNM, whereas PI16^-^/POSTN^high^ fibroblasts varied substantially between patients (figure 3 B-C).

The immune compartment also differed in its cellular composition, with greater representation of myeloid subsets in SSc-VNM and lymphoid populations in a-VNM (supplementary figure S9). CD163^high^/HLA-DR^+^ macrophages were the most abundant immune population in SSc-VNM (mean 23.63%, compared with 8.73% in a-VNM and 11.88% in i-VNM), accompanied by HLA-DR^high^ and CD86^high^ monocytes. a-VNM showed comparatively more HLA-DR^-^ macrophages and CD8^+^ T cells, while i-VNM showed enrichment of CD86^high^ monocytes (supplementary figures S8B and S9B).

### Cellular neighborhoods and spatial interactions

Ten cellular neighborhoods (CNs) captured recurrent myocardial microenvironments (figure 4A and supplementary figure S10). Five CNs were organized around stromal populations: PI16^high^/POSTN^high^ fibroblasts, PI16^-^/POSTN^high^ fibroblasts, myofibroblasts, SOX9^+^ fibroblasts, and CD34^+^ stromal cells. Three CNs were vascular, one was cardiomyocyte-rich and one remained undefined. Compartment-resolved plots showed that each stromal CN incorporated a characteristic combination of immune, vascular and fibroblast populations (figure 4A and supplementary figure S10).

**Figure 4.**
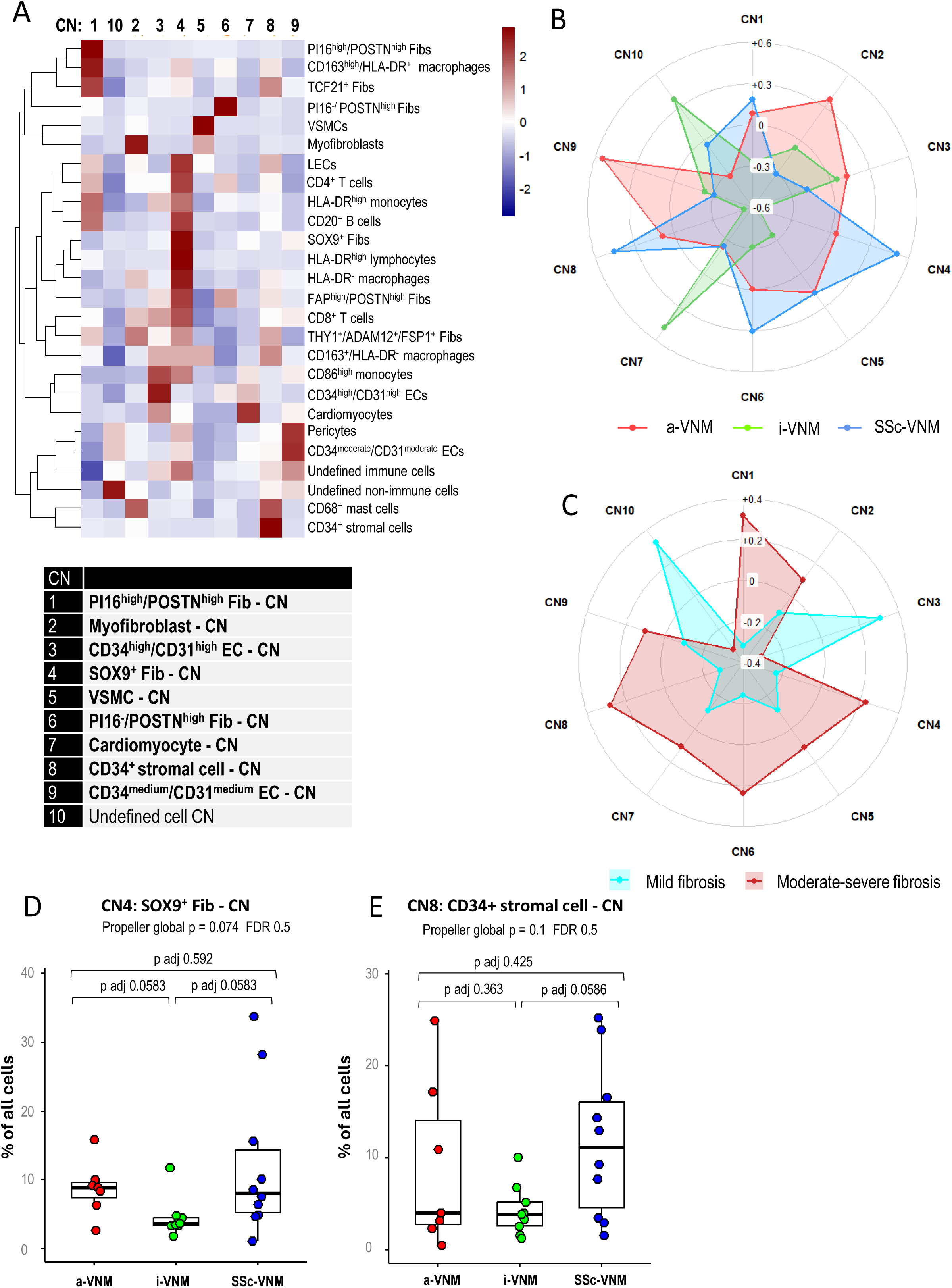
Cellular neighborhoods across disease and fibrosis groups. (A) Cell-type enrichment across 10 cellular neighborhoods (CNs), z-scaled within cell type and hierarchically clustered. (B) Disease-balanced z-scaled CN frequencies in a-VNM, i-VNM and SSc-VNM. (C) Z-scaled CN frequencies in mild (<10% ECM; n=8) versus moderate/severe fibrosis (≥10% ECM; n=14); samples without fibrosis classification were excluded. (D, E) Patient-level frequency of (D) CN4, the SOX9^+^ fibroblast CN, and (E) CN8, the CD34^+^ stromal-cell CN. Propeller global p values, FDR values and Benjamini-Hochberg-adjusted pairwise p values are shown. ECM, extracellular matrix; a-VNM, virus-negative myocarditis associated with a non-SSc systemic autoimmune rheumatic disease; i-VNM, isolated virus-negative myocarditis; SSc-VNM, systemic sclerosis-associated virus-negative myocarditis.

The PI16^high^/POSTN^high^ CN included CD163^high^/HLA-DR^+^ macrophages, TCF21^+^ fibroblasts, HLA-DR^high^ monocytes and CD20^+^ B cells. The myofibroblast CN incorporated THY1^+^/ADAM12^+^/FSP1^+^ fibroblasts, HLA-DR^-^ macrophages, CD8^+^ T cells and CD68^+^ mast cells. The SOX9^+^ CN was particularly enriched in immune cells and included CD20^+^ B cells, monocytes, macrophages and LECs. The CD34^+^ stromal-cell CN combined TCF21^+^ and THY1^+^/ADAM12^+^/FSP1^+^ fibroblasts with CD163^+^/HLA-DR^-^ macrophages and CD68^+^ mast cells. These compositions demonstrate that phenotypically defined fibroblasts/stromal cells reside in reproducible multicellular contexts rather than isolated tissue compartments (figure 4A and supplementary figure S10).

CN frequencies also displayed distinct disease patterns (figure 4B, D-E and supplementary figure S10). SSc-VNM showed higher mean assignment to the SOX9^+^ fibroblast CN (12.1% of all cells versus 8.8% in a-VNM and 4.5% in i-VNM) and the CD34^+^ stromal-cell CN (11.8% versus 9.0% and 4.3%, respectively) (supplementary figure S11).

Interaction analysis further resolved shared and disease-associated cellular networks (figure 5A, supplementary figure S12). Across disease groups, CD34^+^ stromal cells interacted with TCF21^+^ and POSTN^high^ fibroblast subpopulations, and PI16^-^/POSTN^high^ fibroblasts interacted with FAP^high^/POSTN^high^ and PI16^high^/POSTN^high^ fibroblasts. The SSc-VNM cellular networks additionally linked CD34^+^ stromal cells with THY1^+^/ADAM12^+^/FSP1^+^ fibroblasts and linked myofibroblasts with PI16^high^/POSTN^high^ and FAP^high^/POSTN^high^ fibroblasts (supplementary figure S12A).

**Figure 5.**
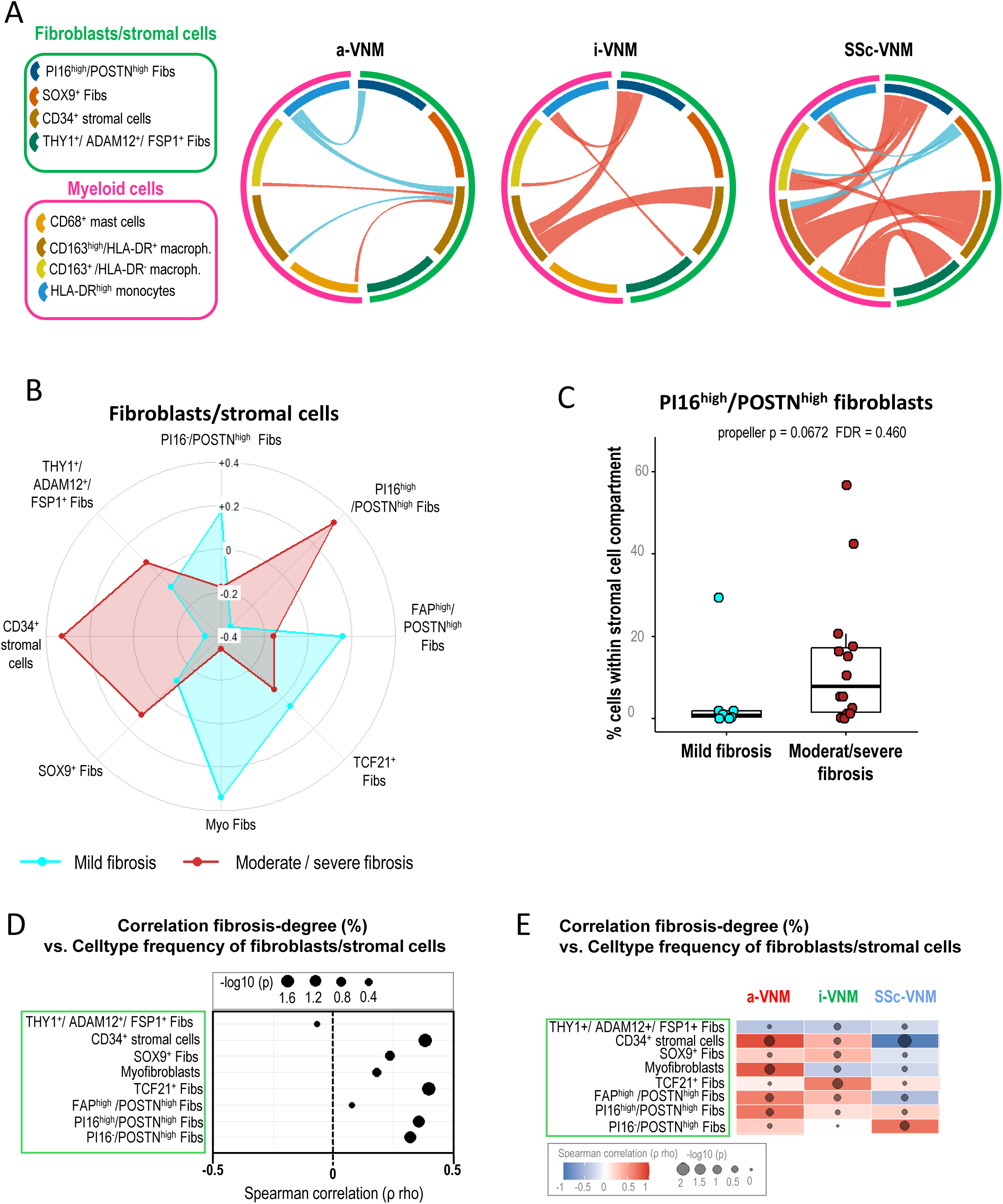
Spatial cell-cell interaction networks across disease groups and distribution of fibroblasts/stromal cells in myocarditis with mild or moderate/severe histological fibrosis. (A) Chord diagrams showing significant permutation-based interactions between selected fibroblast/stromal cell substypes and myeloid cell subsets across myocarditis groups using four-nearest-neighbor graphs. Bidirectional interaction scores were averaged per sample and summed within disease group. Red denotes spatial enrichment and blue spatial avoidance; chord width is proportional to the absolute summed interaction score. (B) Z-scaled stromal phenotypes in mild (<10% ECM; n=8) versus moderate/severe fibrosis (≥10% ECM; n=14), independent of disease group. (C) Patient-level proportions of PI16^high^/POSTN^high^ fibroblasts. (D) Exploratory Spearman correlations between continuous ECM percentage and stromal-phenotype frequency in myocarditis. The x-axis shows Spearman rho, bubble size/position on the y-axis reflects −log10(p). (E) Heatmap illustrating the disease-specific Spearman rho and p values for the correlations between the histological severity of myocardial fibrosis and the frequency of the distinct fibroblast/stromal cell subsets in a-VNM (n=6), i-VNM (n=9) and SSc-VNM (n=7). a-VNM, virus-negative myocarditis associated with a non-SSc systemic autoimmune rheumatic disease; i-VNM, isolated virus-negative myocarditis; SSc-VNM, systemic sclerosis-associated virus-negative myocarditis.

Stromal-immune networks in SSc-VNM converged on CD163^high^/HLA-DR^+^ macrophages and CD68^+^ mast cells. CD34^+^ stromal cells, THY1^+^/ADAM12^+^/FSP1^+^ fibroblasts and PI16^high^/POSTN^high^ fibroblasts each displayed enriched interactions with one or both populations (figure 5A).

### Associations with histological fibrosis and collagen spatial distribution in IMC

The mean fibrotic area was higher in SSc-VNM (16.6%), than in a-VNM (13.6%) and i-VNM (7.2%). Biopsies with ≥10% fibrotic burden showed expansion across much of the fibroblast/stromal compartment compared with those with <10% fibrotic remodeling (figure 5B and supplementary figure S13). PI16^high^/POSTN^high^ fibroblasts were numerically more frequent in biopsies with ≥10% ECM deposition (figure 5C).

Exploratory analyses with the ECM area as a continuous variable revealed disease-specific differences (figure 5E). In the pooled analysis, the extent of the ECM demonstrated positive Spearman correlation coefficients for PI16^high^/POSTN^high^ fibroblasts, PI16^-^/POSTN^high^ fibroblasts and CD34^+^ stromal cells (figure 5D). For myofibroblasts, the clearest positive relationship with ECM area was observed in a-VNM (figure 5E).

Exploratory Spearman analyses assessing potential associations of the continuous fibrosis burden with immune-cell frequencies suggested additional disease-specific associations (supplementary figure S14). In SSc-VNM, CD163^high^/HLA-DR^+^ macrophages showed a positive association with histological fibrosis; in a-VNM, the positive association was most apparent for CD4^+^ T cells; and in i-VNM, positive associations were observed across several immune-cell populations (supplementary figure S14).

IMC-based analysis of spatial relationships of different cell types with collagen in myocarditis demonstrated that the immediate microenvironment (within a radius of 20 µm around the cell) of PI16^high^/POSTN^high^ fibroblasts has the highest median collagen thickness compared with the other fibroblast/stromal cell subsets. Consistently, PI16^high^/POSTN^high^ fibroblasts had the lowest distance to collagen compared with other fibroblast/stromal cell types (figure 6 and supplementary figure S15).

**Figure 6.**
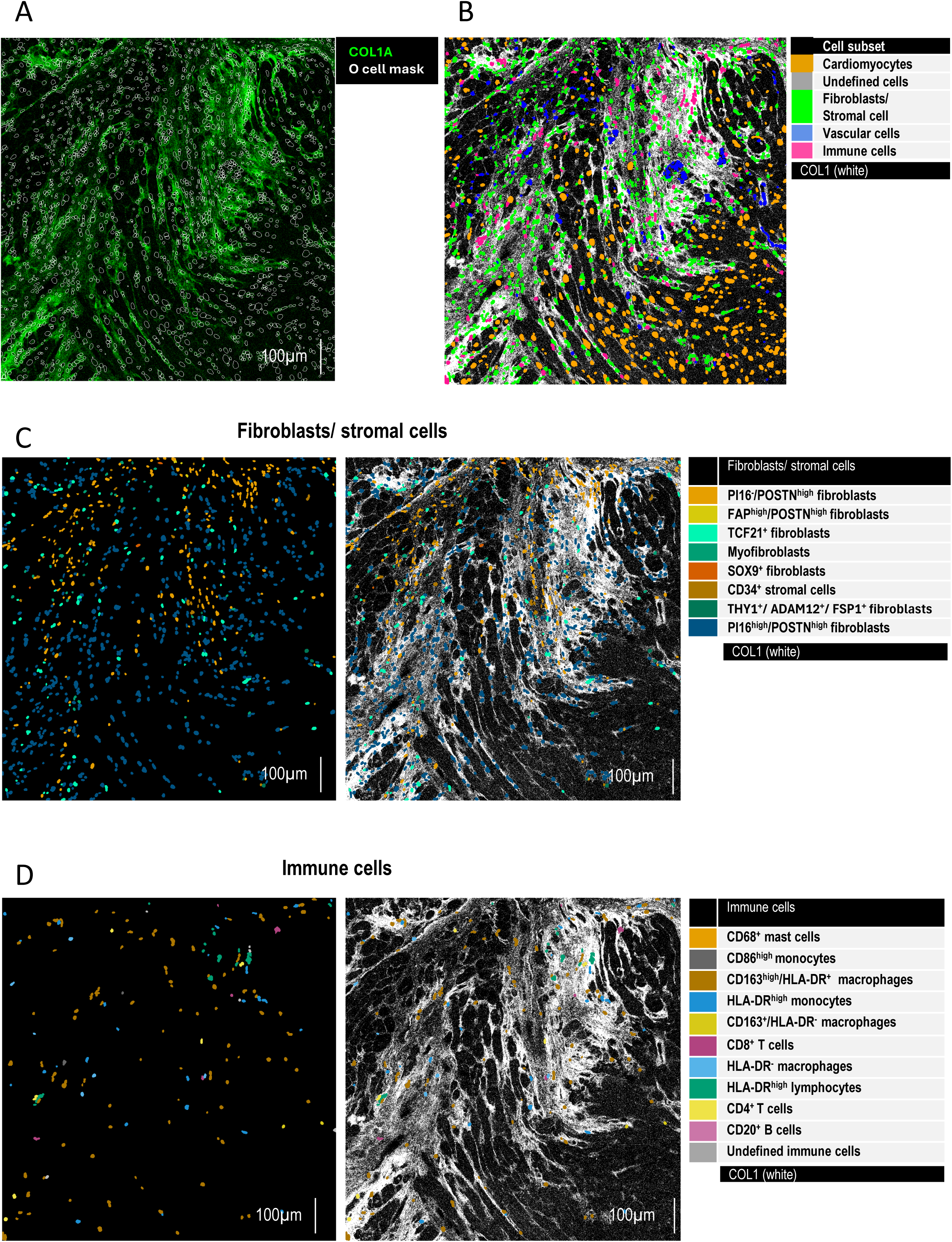
Spatial distribution of fibroblast/stromal phenotypes in relationship to myocardial fibrosis in highly fibrotic VNM. (A) Overlay of collagen signal and cell masks in representative highly fibrotic SSc-VNM, respectively, with ≥ 10% fibrotic remodeling. (B) Corresponding cytomapper image illustrating the main cell subsets: cardiomyocytes in orange, stromal cells in green, vascular cells in blue, and immune cells in pink. (C), Corresponding cytomapper images showing the spatial distribution of the distinct fibroblast/stromal cells clusters (left) and overlay with collagen in white (right) (D) Corresponding cytomapper images showing the spatial distribution of the distinct immune cells clusters (left) and overlay with collagen in white (right). a-VNM, virus-negative myocarditis associated with a non-SSc systemic autoimmune rheumatic disease; i-VNM, isolated virus-negative myocarditis; SSc-VNM, systemic sclerosis-associated virus-negative myocarditis.

Of note, in SSc-VNM CD34^high^/CD31^high^ and CD34^moderate^/CD31^moderate^ ECs demonstrated a significantly higher median collagen thickness within 20 µm compared with a-VNM and i-VNM, suggesting a more pronounced perivascular fibrosis in SSc-VNM (supplementary figure S16). Collectively, these findings support increasing stromal-cell density with greater fibrosis and nominate specific stromal cell and immune-cell populations for future validation.

## Discussion

This study provides a single-cell spatial proteomic map of stromal, vascular and immune organization in SSc-VNM relative to autoimmune non-SSc and isolated virus-negative myocarditis. We demonstrate expansion of the fibroblast/stromal compartment in SSc-VNM with profound changes in the composition of stromal cells/fibroblasts due to numerical increases in CD34^+^ stromal cells and SOX9^+^ and PI16^high^/POSTN^high^ fibroblasts and de-enrichment of FAP^high^/POSTN^high^ fibroblasts and myofibroblasts compared to a-VNM and i-VNM. These changes occurred despite comparable overall proportions of immune cells and vascular cells, highlighting SSc-specific changes of the fibroblast/stromal compartment.

The cardiac stromal map extends single-cell studies showing that human fibrotic remodeling is distributed across multiple fibroblast states.[2–5,20] A prominent SSc-VNM feature was the numerical expansion of CD34^+^ stromal cells, which formed the largest stromal population and contributed to a spatially distinct neighborhood containing TCF21^+^ and THY1^+^/ADAM12^+^/FSP1^+^ fibroblasts, CD163^+^/HLA-DR^-^ macrophages and CD68^+^ mast cells. CD34^+^ stromal cells have been proposed as plastic interstitial populations with progenitor-like properties and the capacity to acquire activated mesenchymal phenotypes during tissue repair.[22,23] Their abundance and integration into macrophage-rich niches support a model in which CD34^+^ stromal cells participate in the organization of chronic myocardial remodeling.

The macrophage findings add another convergent component to this program. Profibrotic macrophage states are established in SSc skin and lung, including spatially organized macrophage-fibroblast signaling.[24–27] CD163^high^/HLA-DR^+^ macrophage counts were increased in SSc-VNM and demonstrated pleiotropic interactions with CD34^+^, PI16^high^/POSTN^high^ and THY1^+^/ADAM12^+^/FSP1^+^ stromal populations. These findings are consistent with a disease-associated stromal-myeloid axis and are in accordance with cardiac single-cell studies that localize macrophage states to activated fibroblast niches after injury, while highlighting that macrophage phenotype and abundance differ across acute injury and chronic heart failure.[2,3] Targeted transcriptomic profiling and *ex-vivo* perturbation will be required to distinguish an SSc-associated macrophage program from treatment exposure or differences in myocarditis phase.

SOX9^+^ fibroblasts defined a complementary spatial state. SOX9 regulates mesenchymal activation and fibrosis across organs and in experimental cardiac injury, where fibroblast-specific Sox9 deletion reduces scar formation and inflammatory recruitment.[28–31] In our cohort, SOX9^+^ fibroblasts were numerically enriched in SSc-VNM and formed an immune-rich neighborhood with CD20^+^ B cells, monocytic cells and lymphatic endothelial cells. B-cell-containing niches are particularly relevant because B-cell activation and tissue infiltration are linked to SSc skin and lung progression and novel B-cell depleting strategies have demonstrated positive effects on skin and lung involvement in therapy-refractory SSc patients.[32–36] These findings provide evidence that SOX9^+^ fibroblasts may organize or respond to a B-cell-containing inflammatory microenvironment. This hypothesis, however, requires verification by functional experiments.

POSTN^high^ fibroblasts represent another major axis of heterogeneity. Three POSTN^high^ states were detected in all disease groups. PI16^high^/POSTN^high^ and PI16^-^/POSTN^high^ fibroblasts showed the strongest intracellular COL1 signal and PI16^high^/POSTN^high^ fibroblasts located in collagen-rich areas. This aligns with experimental lineage and ablation studies and with human spatial data showing that periostin marks activated fibroblast programs after cardiac injury.[4,37–39] FAP^high^/POSTN^high^ fibroblast counts were reduced in SSc-VNM, while PI16^high^/POSTN^high^ fibroblasts were numerically increased compared with a-VNM and i-VNM and showed positive associations with the severity of histological fibrosis. These three populations demonstrate differences in type I collagen expression, are located within distinct neighborhoods and have different interaction partners, all of which supports functional separation of POSTN^high^ states rather than a single fibroblast subset. PI16 expression provides an additional link to cross-tissue fibroblast organization. PI16^+^ fibroblasts are conserved in human and mouse atlases and are frequently positioned as adventitial or reservoir-like populations capable of generating more specialized activated states.[40–42] The coexistence of PI16^high^/POSTN^high^ and PI16^-^/POSTN^high^ fibroblasts in myocarditis is compatible with a continuum of activation, but the current protein panel and cross-sectional design cannot establish a developmental trajectory.

The lower proportion of α-SMA^high^ myofibroblasts in SSc-VNM despite greater stromal representation and more extensive histological fibrosis suggests that α-SMA^high^ myofibroblasts are not the main matrix-producing stromal population in the context of SSc-VNM. Indeed, the strongest type I collagen expression was observed in POSTN^high^ fibroblasts. This is consistent with recent transcriptomic analysis of SSc-pHI, which identified COL1^high^/POSTN^high^ fibroblasts as a distinct fibroblast population separate from myofibroblasts.[21] Together, these findings indicate that POSTN^high^ fibroblasts, rather than α-SMA^high^ myofibroblasts, are the dominant source of accumulating type I collagen in SSc-VNM. This concept is further supported by emerging human cardiac data demonstrating substantial extracellular-matrix production by non-myofibroblast stromal cell populations.[4,5,20]

The TPSAB1^+^/CD68^+^ mast-cell phenotype also warrants further investigation. The two antibody channels were separated by 60 mass units, making conventional 16-mass-unit oxide spillover unlikely. In addition, double-positive cells were clearly detectable in spatially resolved images. Moreover, a related protein-defined mast-cell population has previously been described in colorectal-cancer stroma.[19] In SSc-VNM, these cells were numerically enriched and localized particularly to CD34^+^ stromal and myofibroblast neighborhoods. However, they remain a relatively rare cell population. Their low frequencies, eventually together with loss of these cells during dissociation and post-translational regulation may explain why these cells have not been recovered by conventional scRNASeq approaches. Further validation with orthogonal co-staining with additional mast-cell and myeloid markers are required before assigning a distinct activated-mast-cell program.

Limitations of our study include the small retrospective cohort, lack of a non-inflamed control group, and incomplete clinical characterization. Unlike patients in the two myocarditis control groups, most patients with SSc-VNM received immunosuppressive therapies, which may have influenced immune cell composition and activation states. Furthermore, the three myocarditis groups were not balanced for further potential confounders such as sex, age, and phase of myocarditis. Endomyocardial biopsy sampling may not fully capture diffuse myocardial heterogeneity, and annotation with a limited panel of protein markers may not fully resolve each functional state. These considerations are especially important in inflammatory cardiomyopathy, where immune composition and repair programs vary with disease phase.[43,44] However, this study is the first to provide spatially resolved multiplexed protein-based phenotyping of different VNM entities at single cell resolution and a comparison of SSc-VNM with two myocarditis control groups. The study also links cell abundance, neighborhood composition, pairwise interactions and histological fibrosis within the same biopsies.

In conclusion, SSc-VNM shows reproducible expansion and spatial redistribution of the cardiac stromal compartment and a distinct balance of FAP^high^/POSTN^high^ fibroblasts and myofibroblasts. The accompanying spatial map demonstrates increased abundance of CD34^+^, SOX9^+^ and POSTN^+^ stromal cells and reorganization of the cellular niches with enrichment of macrophages, mast-cells, and B-cells within the local microenvironments. This study provides both first protein-based disease mapping and identifies cellular candidates for future mechanistic studies.

## Declarations

### Data availability statement

Data are available from the corresponding authors on reasonable request, subject to ethical and institutional restrictions.

### Patient and public involvement

Patients and/or the public were not involved in the design, conduct, reporting or dissemination plans of this retrospective tissue study.

### Ethics approval

All participants provided written informed consent. The study was approved by the responsible institutional review boards of the Medical Faculty of Heinrich Heine University Düsseldorf, Vita-Salute San Raffaele University Milan and University Hospital Erlangen.

### Funding

The project was supported by the following grants: Grants DI 1537/17-1, DI 1537/20-1, DI 1537/22-1, DI 1537/23-1, DI 1537/27-1, and DI 1537/28-1 of the German Research Foundation (JHWD), MA 9219/2-1 of the German Research Foundation (AEM), 493659010 of the German Research Foundation, SFB CRC1181 (project C01) of the German Research Foundation, CRC1774 (project B03, project number 556052979), grants 2021_EKEA.03 (AEM) and 2022_EKMS.02 (AEM) of the Else-Kröner-Fresenius-Foundation, The Edith Busch and World Scleroderma Foundation Research Grant Programme 2022-2023 (AEM), the Research Committee of the Medical Faculty of the Heinrich-Heine University Düsseldorf (Forschungskommission; ID 2022-18, ID 2023-33 and ID 2023-31 to AHG, AEM, and JHWD, respectively), an unrestricted research grant from the Hiller-Foundation (JHWD), and a Career Support Award of Medicine of the Ernst Jung Foundation (JHWD).

### Contributors

**Conceptualization**: JHWD, AHG.

**Data curation**: ANS, AEM, YNL, TF.

**Formal analysis**: ANS, AEM, YNL, TF.

**Funding acquisition**: AEM, JHWD, AHG.

**Investigation**: ANS, AEM, YNL, TF, SR, MDG, CB, AHG.

**Methodology**: ANS, AEM, YNL, TF.

**Project administration**: JHWD, AHG.

**Resources**: all authors.

**Software**: ANS, AEM, YNL, TF.

**Supervision**: JHWD, AHG.

**Validation**: all authors.

**Visualization**: ANS, JHWD, AHG.

**Writing – original draft**: ANS, JHWD, AHG.

**Writing – review and editing**: all authors.

## Acknowledgements

We thank Christopher Liebel, Lukas Sokolowski and Philipp Steinbrecher for excellent technical support.

## Competing interests

EZ reports speaker honoraria from Abiomed (Johnson & Johnson) and travel support from Bayer and Pfizer. JHWD reports grants or contracts from AbbVie, Agomab, Anamar, argenx, ARXX, Bristol Myers Squibb, Bayer, Boehringer Ingelheim, Candid, Cantargia, Celgene, CSL Behring, Eli Lilly, ExoTherapeutics, GSK, Lassen, Kyverna, Mestag, Novartis, Otsuka, Sanofi, Redx and UCB; consulting fees from AbbVie, Anamar, ARXX, AstraZeneca, Boehringer Ingelheim, Celgene, Genentech/Roche, GSK, Novartis, Pfizer, Roche, Sulis, Spica, Tyra, UCB and Xencor; honoraria from AbbVie, Anamar, ARXX/Caluna, AstraZeneca, Boehringer Ingelheim, Eli Lilly, Genentech/Roche, GSK, Merck, Novartis and UCB; expert-testimony payments from AbbVie, Anamar, ARXX/Caluna, AstraZeneca, Bayer, Boehringer Ingelheim, Cantargia, Genentech, Quell, Kyverna, Novartis, Prolium, Roche and UCB; meeting or travel support from AbbVie and SOBI; and stock or stock options in 4D Science and FibroCure. MB-H: honoraria for lectures from Novartis, Pfizer, Sanofi.

## References

1 Frangogiannis NG. Cardiac fibrosis. Cardiovasc Res. 2021;117:1450–88. doi: 10.1093/cvr/cvaa324

2 Kuppe C, Ramirez Flores RO, Li Z, et al. Spatial multi-omic map of human myocardial infarction. Nature. 2022;608:766–77. doi: 10.1038/s41586-022-05060-x

3 Koenig AL, Shchukina I, Amrute J, et al. Single-cell transcriptomics reveals cell-type-specific diversification in human heart failure. Nat Cardiovasc Res. 2022;1:263–80. doi: 10.1038/s44161-022-00028-6

4 Amrute JM, Luo X, Penna V, et al. Targeting immune-fibroblast cell communication in heart failure. Nature. 2024;635:423–33. doi: 10.1038/s41586-024-08008-5

5 Youness M, Ekhteraei-Tousi S, Nagaraju CK, et al. Location and aetiology are determinants of fibroblast activation and heterogeneity in the failing human heart. Genome Med. 2025;17:155. doi: 10.1186/s13073-025-01580-z

6 Denton CP, Khanna D. Systemic sclerosis. Lancet Lond Engl. 2017;390:1685–99. doi: 10.1016/S0140-6736(17)30933-9

7 Elhai M, Meune C, Boubaya M, et al. Mapping and predicting mortality from systemic sclerosis. Ann Rheum Dis. 2017;76:1897–905. doi: 10.1136/annrheumdis-2017-211448

8 Bruni C, Buch MH, Furst DE, et al. Primary systemic sclerosis heart involvement: A systematic literature review and preliminary data-driven, consensus-based WSF/HFA definition. J Scleroderma Relat Disord. 2022;7:24–32. doi: 10.1177/23971983211053246

9 Györfi A-H, Filla T, Polzin A, et al. Evaluation of Systemic Sclerosis Primary Heart Involvement and Chronic Heart Failure in the European Scleroderma Trials and Research Cohort. J Am Heart Assoc. 2025;14:e036730. doi: 10.1161/JAHA.124.036730

10 Mueller KAL, Mueller II, Eppler D, et al. Clinical and histopathological features of patients with systemic sclerosis undergoing endomyocardial biopsy. PloS One. 2015;10:e0126707. doi: 10.1371/journal.pone.0126707

11 De Luca G, Campochiaro C, De Santis M, et al. Systemic sclerosis myocarditis has unique clinical, histological and prognostic features: a comparative histological analysis. Rheumatol Oxf Engl. 2020;59:2523–33. doi: 10.1093/rheumatology/kez658

12 Rius Rigau A, Liang M, Devakumar V, et al. Imaging mass cytometry-based characterisation of fibroblast subsets and their cellular niches in systemic sclerosis. Ann Rheum Dis. 2024;ard-2024-226336. doi: 10.1136/ard-2024-226336

13 Li Y-N, Filla T, Györfi A-H, et al. Spatially informed phenotyping by cyclic-in-situ-hybridisation identifies novel fibroblast populations and their pathogenic niches in systemic sclerosis. Ann Rheum Dis. 2025;84:1852–64. doi: 10.1016/j.ard.2025.06.002

14 Valenzi E, Bulik M, Tabib T, et al. Single-cell analysis reveals fibroblast heterogeneity and myofibroblasts in systemic sclerosis-associated interstitial lung disease. Ann Rheum Dis. 2019;78:1379–87. doi: 10.1136/annrheumdis-2018-214865

15 Isomura Y, Nakano Y, Kubota Y, et al. Transcriptomic profiling of myocardial biopsies reveals metabolic suppression and autophagy activation in systemic sclerosis-associated primary heart involvement. J Scleroderma Relat Disord. 2026;11:e000003. doi: 10.1136/jsrd-2026-000003

16 Aretz HT, Billingham ME, Edwards WD, et al. Myocarditis. A histopathologic definition and classification. Am J Cardiovasc Pathol. 1987;1:3–14.

17 van den Hoogen F, Khanna D, Fransen J, et al. 2013 classification criteria for systemic sclerosis: an American college of rheumatology/European league against rheumatism collaborative initiative. Ann Rheum Dis. 2013;72:1747–55. doi: 10.1136/annrheumdis-2013-204424

18 Windhager J, Zanotelli VRT, Schulz D, et al. An end-to-end workflow for multiplexed image processing and analysis. Nat Protoc. 2023;18:3565–613. doi: 10.1038/s41596-023-00881-0

19 Arelaki S, Arampatzioglou A, Kambas K, et al. Mast cells co-expressing CD68 and inorganic polyphosphate are linked with colorectal cancer. PloS One. 2018;13:e0193089. doi: 10.1371/journal.pone.0193089

20 Van Linthout S, Matz I, González A, et al. Cardiac fibroblasts in myocardial injury and heart failure. Eur Heart J. 2026;47:2271–85. doi: 10.1093/eurheartj/ehaf902

21 Micu A, Matei A-E, Li Y-N, et al. OC.46 Spatial transcriptomic-based phenotyping of the fibroblast niches in systemic sclerosis-associated primary heart involvement. Oral Communications. BMJ Publishing Group Ltd 2026:A48–9.

22 Díaz-Flores L, Gutiérrez R, García MP, et al. CD34+ stromal cells/fibroblasts/fibrocytes/telocytes as a tissue reserve and a principal source of mesenchymal cells. Location, morphology, function and role in pathology. Histol Histopathol. 2014;29:831–70. doi: 10.14670/HH-29.831

23 Díaz-Flores L, Gutiérrez R, García MP, et al. Human resident CD34+ stromal cells/telocytes have progenitor capacity and are a source of αSMA+ cells during repair. Histol Histopathol. 2015;30:615–27. doi: 10.14670/HH-30.615

24 Xue D, Tabib T, Morse C, et al. Expansion of Fcγ Receptor IIIa-Positive Macrophages, Ficolin 1-Positive Monocyte-Derived Dendritic Cells, and Plasmacytoid Dendritic Cells Associated With Severe Skin Disease in Systemic Sclerosis. Arthritis Rheumatol Hoboken NJ. 2022;74:329–41. doi: 10.1002/art.41813

25 Jarnagin HC, Parvizi R, Gong Z, et al. Multimodal analyses of early, untreated systemic sclerosis skin identify a proinflammatory vascular niche of macrophage-fibroblast signaling. JCI Insight. 2026;11:e198954. doi: 10.1172/jci.insight.198954

26 Papazoglou A, Huang M, Bulik M, et al. Epigenetic Regulation of Profibrotic Macrophages in Systemic Sclerosis-Associated Interstitial Lung Disease. Arthritis Rheumatol Hoboken NJ. 2022;74:2003–14. doi: 10.1002/art.42286

27 Gotelli E, Soldano S, Feghali-Bostwick C, et al. Prevalence of hybrid TLR4+M2 monocytes/macrophages in peripheral blood and lung of systemic sclerosis patients with interstitial lung disease. Front Immunol. 2024;15:1488867. doi: 10.3389/fimmu.2024.1488867

28 Hanley KP, Oakley F, Sugden S, et al. Ectopic SOX9 mediates extracellular matrix deposition characteristic of organ fibrosis. J Biol Chem. 2008;283:14063–71. doi: 10.1074/jbc.M707390200

29 Lacraz GPA, Junker JP, Gladka MM, et al. Tomo-Seq Identifies SOX9 as a Key Regulator of Cardiac Fibrosis During Ischemic Injury. Circulation. 2017;136:1396–409. doi: 10.1161/CIRCULATIONAHA.117.027832

30 Scharf GM, Kilian K, Cordero J, et al. Inactivation of Sox9 in fibroblasts reduces cardiac fibrosis and inflammation. JCI Insight. 2019;5:e126721. doi: 10.1172/jci.insight.126721

31 Raza S, Jokl E, Pritchett J, et al. SOX9 is required for kidney fibrosis and activates NAV3 to drive renal myofibroblast function. Sci Signal. 2021;14:eabb4282. doi: 10.1126/scisignal.abb4282

32 Bosello S, Angelucci C, Lama G, et al. Characterization of inflammatory cell infiltrate of scleroderma skin: B cells and skin score progression. Arthritis Res Ther. 2018;20:75. doi: 10.1186/s13075-018-1569-0

33 Lafyatis R, O’Hara C, Feghali-Bostwick CA, et al. B cell infiltration in systemic sclerosis-associated interstitial lung disease. Arthritis Rheum. 2007;56:3167–8. doi: 10.1002/art.22847

34 Jia G, Ramalingam TR, Heiden JV, et al. An interleukin 6 responsive plasma cell signature is associated with disease progression in systemic sclerosis interstitial lung disease. iScience. 2023;26:108133. doi: 10.1016/j.isci.2023.108133

35 Düsing C, Györfi A-H, Stütz AN, et al. Bispecific T cell engagers for treatment-refractory autoimmune connective tissue diseases. Nat Med. 2026;32:1530–42. doi: 10.1038/s41591-026-04238-4

36 Auth J, Müller F, Völkl S, et al. CD19-targeting CAR T-cell therapy in patients with diffuse systemic sclerosis: a case series. Lancet Rheumatol. 2025;7:e83–e93. doi: 10.1016/S2665-9913(24)00282-0

37 Shimazaki M, Nakamura K, Kii I, et al. Periostin is essential for cardiac healing after acute myocardial infarction. J Exp Med. 2008;205:295–303. doi: 10.1084/jem.20071297

38 Kaur H, Takefuji M, Ngai CY, et al. Targeted Ablation of Periostin-Expressing Activated Fibroblasts Prevents Adverse Cardiac Remodeling in Mice. Circ Res. 2016;118:1906–17. doi: 10.1161/CIRCRESAHA.116.308643

39 Kanisicak O, Khalil H, Ivey MJ, et al. Genetic lineage tracing defines myofibroblast origin and function in the injured heart. Nat Commun. 2016;7:12260. doi: 10.1038/ncomms12260

40 Buechler MB, Pradhan RN, Krishnamurty AT, et al. Cross-tissue organization of the fibroblast lineage. Nature. 2021;593:575–9. doi: 10.1038/s41586-021-03549-5

41 Gao Y, Li J, Cheng W, et al. Cross-tissue human fibroblast atlas reveals myofibroblast subtypes with distinct roles in immune modulation. Cancer Cell. 2024;42:1764–1783.e10. doi: 10.1016/j.ccell.2024.08.020

42 Patrick R, Janbandhu V, Tallapragada V, et al. Integration mapping of cardiac fibroblast single-cell transcriptomes elucidates cellular principles of fibrosis in diverse pathologies. Sci Adv. 2024;10:eadk8501. doi: 10.1126/sciadv.adk8501

43 Tschöpe C, Ammirati E, Bozkurt B, et al. Myocarditis and inflammatory cardiomyopathy: current evidence and future directions. Nat Rev Cardiol. 2021;18:169–93. doi: 10.1038/s41569-020-00435-x

44 Chen R, Zhang H, Tang B, et al. Macrophages in cardiovascular diseases: molecular mechanisms and therapeutic targets. Signal Transduct Target Ther. 2024;9:130. doi: 10.1038/s41392-024-01840-1

45 Leone O, Veinot JP, Angelini A, et al. 2011 consensus statement on endomyocardial biopsy from the Association for European Cardiovascular Pathology and the Society for Cardiovascular Pathology. Cardiovasc Pathol Off J Soc Cardiovasc Pathol. 2012;21:245–74. doi: 10.1016/j.carpath.2011.10.001

46 Rius Rigau A, Li Y-N, Matei A-E, et al. Characterization of Vascular Niche in Systemic Sclerosis by Spatial Proteomics. Circ Res. 2024;134:875–91. doi: 10.1161/CIRCRESAHA.123.323299

47 Grönberg C, Rattik S, Tran-Manh C, et al. Combined inhibition of IL-1, IL-33 and IL-36 signalling by targeting IL1RAP ameliorates skin and lung fibrosis in preclinical models of systemic sclerosis. Ann Rheum Dis. 2024;83:1156–68. doi: 10.1136/ard-2023-225158

48 Györfi A-H, Matei A-E, Fuchs M, et al. Engrailed 1 coordinates cytoskeletal reorganization to induce myofibroblast differentiation. J Exp Med. 2021;218:e20201916. doi: 10.1084/jem.20201916

