## Supplementary material for "Spatial proteomic profiling of cardiac stromal populations and microenvironments in systemic sclerosis-associated primary myocardial involvement": Suppl.tables

**Supplementary table S1. Patient characteristics**

The numbers in brackets for SSc patients show the respective modified Rodnan skin score (mRSS). Abbreviations: a-VNM, autoimmune rheumatic disease-associated virus-negative myocarditis; EMB, endomyocardial biopsy; i-VNM, isolated virus-negative myocarditis; n/a, not available; SSc, systemic sclerosis; SSc-VNM, systemic sclerosis-associated virus-negative myocarditis, SARD, systemic autoimmune rheumatic disease; ABs,autoantibodies; Histopath., histopathological; Immunmod: immunomodulatory; LVEF, left ventricular ejection fraction; CMR, cardiac magnetic resonance; TTE, transthoracic echocardiography; hsTnT, high sensitive Troponin T (in brackets upper limit of normal); ASyS, anti-synthetase syndrome; UCTD, undifferentiated connective tissue disease; SLE, systemic lupus erythematosus; ILD, interstitial lung disease; GERD, gastroesophageal reflux disease; DU, digital ulcers; RP, Raynaud’s phenomenon; SVT, supraventricular tachycardia; ICD, implantable cardioverter-defibrillator; VEBs, ventricular ectopic beats; NSVT, non-sustained ventricular tachycardia; SVEBs, supraventricular ectopic beats; VF, ventricular fibrillation; AVB, atrioventricular block; AT, atrial tachycardia; WPW, Wolff Parkinson White Syndrome; RBBB, right bundle branch block; MMF, mycophenolate mofetil; HCQ, hydroxychloroquine; AZA, azathioprine; MTX, methotrexate; Pred, prednisolone; CMR parameters: T1 and T2 refer to myocardial relaxation times assessed by CMR mapping; LGE, late gadolinium enhancement. Increased native (T1 and T2 values were defined according to local reference ranges); n/a, not available date.

| **Patient-ID** | **Sex** | **Age** | **ABs** | **SARD** | **Histopath. fibrosis (%) in EMB** | **Immunomod. treatment at time of EMB** | **Comorbidities** | **Arrhythmia** | **LVEF in CMR or TTE** | **hsTnT [ng/L]**  **at time of EMB** | **CMR parameters** |
| --- | --- | --- | --- | --- | --- | --- | --- | --- | --- | --- | --- |
| **a-VNM_1** | M | 55 y | PL12 | ASyS | 3,5 | none | hypothyroidism, former smoker | None | 58% | 490 (<14) | edema  T1 increased  T2 increaced  LGE n/a |
| **a-VNM_2** | M | 59 y | Neg. | Sarcoidosis | n/a | none | hyperthyroidism | SVT, ICD | 49% | 19.5 (<14) | n/a |
| **a-VNM_3** | F | 44 y | TIF1γ | UCTD | 16 | none | pituitary hyperplasia, smoker | sinus tachycardia | n/a | 3 (<14) | edema  T1 increased  T2 increaced  LGE present |
| **a-VNM_4** | M | 56 y | EJ1 | UCTD | 15 | none | history of deep venous thrombosis, benign prostatic hyperplasia, former smoker | VEBs | 30% | 12 (<14) | edema n/a  T1 n/a  T2 n/a  LGE present |
| **a-VNM_5** | F | 53 y | PM-Scl100 | UCTD | 30 | none | none | VEBs, NSVTs | 61% | 140 (<14) | minimal edema  T1 n/a  T2 increased  LGE present |
| **a-VNM_6** | M | 77 y | Jo1 | ASyS | 12 | MMF | asthma, dyslipidemia, kidney lithiasis, ischemic heart disease | SVEBs | 56% | 29 (<14) | edema n/a  T1 increased  T2 n/a  LGE present |
| **a-VNM_7** | F | 30 y | dsDNA | SLE | 5 | Pred, HCQ, MMF | vitiligo, former smoker | sinus tachycardia | 65% | 0.65 (<0.034) | edema  T1 n/a  T2 n/a  LGE present |
| **i-VNM_1** | M | 23 y | Neg. | no | 7 | none | psoriasis | former ablation by SVT | 60% | 52 (<14) | edema  T1 n/a  T2 increased  LGE present |
| **i-VNM_2** | M | 74 y | Neg. | no | 10 | none | dyslipidemia, carotid stenosis, recurrent herpes labialis and herpes zoster | NSVT episode | n/a | 228 (<14) | n/a |
| **i-VNM_3** | M | 41 y | Neg. | no | 2 | none | none | ICD implantation after cardiac arrest | 45% | 34 (<14) | edema n/a  T1 n/a  T2 in/a  LGE present |
| **i-VNM_4** | M | 74 y | Neg. | no | 12 | none | arterial hypertension, polycystic kidney diseases, microcytic anemia, previous smoker | none | 68% | 35.7 (<14) | no edema  T1 increased  T2 increased  LGE present |
| **i-VNM_5** | F | 42 y | Neg. | no | 11 | none | none | ICD implantation after cardiac arrest, VF | 66% | n/a | minimal edeam  T1 increased  T2 n/a  LGE absent |
| **i-VNM_6** | F | 43 y | Neg. | no | 9,5 | none | fibromyalgia, hyperhomocysteinemia, ovarian cyst surgery, previous smoker | NSVTs | 69% | 52 (<14) | minimal edema  T1 n/a  T2 n/a  LGE absent |
| **i-VNM_7** | M | 67 y | Neg. | no | 0 | none | arterial hypertension, hyperuricemia, carotid stenosis, previous smoker, former acute myocardial infarction | AVB II°, NSVTs | 59% | 16.9 (<14) | edema  T1 increased  T2 increased  LGE present |
| **i-VNM_8** | M | 51 y | Neg. | no | 4 | none | previous smoker | VEBs | n/a | 39.1 (<14) | n/a |
| **i-VNM_9** | M | 46 y | Neg. | no | 10 | none | GERD, smoker | none | 59% | n/a | edema  T1 increased  T2 increased  LGE present |
| **SSc-VNM_1** | F | 36 y | RNP-U1 | dcSSc  (13) | 5 | Pred, HCQ, AZA | thalassemia minor, chronic multinodular thyroiditis | Incomplete RBBB, early repolarization | n/a | 360 (<14) | n/a |
| **SSc-VNM_2** | F | 63 y | CENP | lcSSc  (4) | 15 | Pred, ciclosporin | impaired fasting blood glucose, osteoporosis, depressive syndrome, fibrocystic mastopathy | polymorphic VEBs, couplets/triplets, SVEBs, AT episodes | 60% | 260 (<14) | edema  T1 increased  T2 increased  LGE n/a |
| **SSc-VNM_3** | F | 32 y | PM-Scl100,  CENP | lcSSc  (8) | 10 | none | hypothyroidism | none | n/a (preserved) | 21 (<14) | edema n/a  T1 n/a  T2 increased  LGE present |
| **SSc-VNM_4** | F | 39 y | CENP | SSc sine scleroderma | 24 | colchicine | none | polymorphic VEBs, couplets | 52% | 3 (<14) | no edema  T1 borderline  T2 borderline  LGE present |
| **SSc-VNM_5** | F | 58 y | Th/To | lcSSc  (n/a) | 10 | none | autoimmune thyroiditis | WPW with recurrent tachycardia | n/a | 74 (<14) | edema n/a  T1 n/a  T2 n/a  LGE present |
| **SSc-VNM_6** | M | 31 y | Scl70 | dcSSc  (n/a) | n/a | MTX | none | VEBs, NSVTs, RBBB | n/a | 156 (<34) | n/a |
| **SSc-VNM_7** | F | 52 y | n/a | n/a | n/a | none | none | none | n/a | 1347 (<34) | n/a |
| **SSc-VNM_8** | M | 58 y | Fibrillarin | dcSSc  (13) | 40 | MMF | arterial hypertension | VEBs, SVEBs | 60% | 107 (<14) | edema  T1 increased  T2 increased  LGE present |
| **SSc-VNM_9** | M | 61 y | Scl70 | dcSSc  (17) | 12 | MMF | none | VEBs, couplets/triplets, SVEBs | 45% | 164 (<14) | edema  T1 n/a  T2 increased  LGE present |
| **SSc-VNM_10** | F | 59 y | RNP-U1 | lcSSc  (0) | n/a | Pred, MMF | arterial hypertension, hypercholesterolemia | RBBB | 65% | n/a | n/a |

**Supplementary table S2. IMC staining antibodies and metal isotopes**

Antibodies marked with an asterisk were purchased preconjugated from Fluidigm/Standard BioTools; all remaining antibodies were conjugated with the Maxpar X8 Antibody Labeling Kit. Working concentrations are stated per 100 µL staining solution.

Companies: SBT, Standard BioTools, South San Francisco, California, USA; Sigma-Aldrich, St Louis, Missouri, USA; Abcam, Cambridge, UK; BioLegend, San Diego, California, USA; Invitrogen, Carlsbad, California, USA; Thermo Fisher Scientific, Waltham, Massachusetts, USA.

| **Metal** | **Company** | **Cat. No.** | **Antigen** | **Antigen Name** | **Clone** | **Company** | **Cat.No.** | **Dilution** |
| --- | --- | --- | --- | --- | --- | --- | --- | --- |
| 89Y | Sigma-Aldrich | 204919-10G | TNNT2 | Troponin T Type 2 (Cardiac) | 1C11 | Abcam | ab8295 | 1:100 |
| 115In | Sigma-Aldrich | 203440-1G | TPSAB1 | Tryptase Alpha/Beta 1 | AA1 | Biolegend | 369402 | 1:600 |
| 141Pr | SBT/ Fluidigm | S00023 | HLA-DR | Major Histocompatibility Complex Class II, DR Beta 1 | LN3 | Invitrogen | 14-9956-82 | 1:400 |
| 143Nd | SBT/ Fluidigm | S00016 | CD34 | CD34 | QBEND/10 | Invitrogen | MA5-18091 | 1:800 |
| 144Nd | ***** | ***** | CD14 | CD14 | EPR3653 | SBT/ Fluidigm | 3144025D | 1:200 |
| 145Nd | SBT/ Fluidigm | S00018 | PECAM1=CD31 | Platelet And Endothelial Cell Adhesion Molecule 1 = CD31 | polyclonal goat | R&D | AF3628 | 1:200 |
| 147Sm | ***** | ***** | CD163 | CD163 Scavenger Receptor | EDHu-1 | SBT/ Fluidigm | 3147021D | 1:200 |
| 148Nd | SBT/ Fluidigm | S00021 | PDGFRA | Platelet Derived Growth Factor Receptor Alpha | polyclonal goat | R&D | AF307NA | 1:200 |
| 150Nd | SBT/ Fluidigm | S00014 | ACTA2  = α-SMA | Actin Alpha 2, Smooth Muscle | 1A4 | Sigma-Aldrich | CBL171-I | 1:200 |
| 151Eu | SBT/ Fluidigm | S00024 | PDGFRB | Platelet Derived Growth Factor Receptor Beta | 18A2 | Biolegend | 323602 | 1:100 |
| 152Sm | ***** | ***** | PTPRC =CD45 | Protein Tyrosine Phosphatase Receptor Type C  =CD45 | D9M8I | SBT/ Fluidigm | 3152018D | 1:200 |
| 153Eu | SBT/ Fluidigm | S00026 | MS4A1 =CD20 | Membrane Spanning 4-Domains A1  =CD20 | H1 | BD | 555677 | 1:800 |
| 154Sm | SBT/ Fluidigm | S00027 | ZEB1 | Zinc Finger E-Box Binding Homeobox 1 | 3G6 | Invitrogen | 14-9741-82 | 1:300 |
| 155Gd | SBT/ Fluidigm | S00098 | CD86 | CD86 | polyclonal  goat | R&D | AF-141-NA | 1:400 |
| 156Gd | ***** | ***** | CD4 | CD4 | EPR6855 | SBT/ Fluidigm | 3156033D | 1:200 |
| 158Gd | SBT/ Fluidigm | S00029 | POSTN | Periostin | polyclonal rabbit | Abcam | ab14041 | 1:200 |
| 159Tb | SBT/ Fluidigm | S00030 | SOX9 | SRY-Box Transcription Factor 9 | E-9 | Santa Cruz | sc-116505X | 1:500 |
| 160Gd | SBT/ Fluidigm | S00031 | PDPN | Podoplanin | LpMab-21 | Biolegend | 395002 | 1:200 |
| 161Dy | SBT/ Fluidigm | S000102 | THY1 | Thy-1 Cell Surface Antigen | EPR3133 | Abcam | ab226123 | 1:600 |
| 162Dy | ***** | ***** | CD8A | CD8A | C8/144B | SBT/ Fluidigm | 3162034D | 1:800 |
| 163Dy | SBT/ Fluidigm | S000104 | COL1 | Collagen Type I Alpha 1 Chain | EPR7785 | Abcam | ab215969 | 1:200 |
| 165Ho | SBT/ Fluidigm | S00034 | VWF | Von Willebrand Factor | polyclonal rabbit | Sigma Aldrich | Ab7356 | 1:400 |
| 166Er | SBT/ Fluidigm | S00035 | ADAM12 | ADAM Metallopeptidase Domain 12 | 1G3 | Novus | H00008038-M01 | 1:400 |
| 169Tm | SBT/ Fluidigm | S00038 | PI16 | Peptidase Inhibitor 16 | polyclonal goat | R&D | AF4980 | 1:500 |
| 170Er | ***** | ***** | CD3 | CD3 | polyclonal C-Terminal | SBT/ Fluidigm | 3170019D | 1:400 |
| 171Yb | SBT/ Fluidigm | S00040 | S100A4  =FSP1 | S100 Calcium Binding Protein A4  = Fibroblast-specific protein 1 | 1F12-1G7 | Novus | N00006275-M01 | 1:600 |
| 172Yb | SBT/ Fluidigm | S00041 | TCF21 | Transcription Factor 21 | EPR13449 | Abcam | ab250578 | 1:400 |
| 174Yb | SBT/ Fluidigm | S00042 | FAP | Fibroblast Activation Protein Alpha | polyclonal sheep | R&D | AF3715 | 1:200 |
| 175Lu | SBT/ Fluidigm | S00043 | CD68 | CD68 | KP1 | Biolegend | 916104 | 1:800 |
| 176Yb | ***** | ***** | Histone 3 | Histone 3 = nuclear marker | D1H2 | SBT/ Fluidigm | 4276023D | 1:1600 |
| 191Ir  /193Ir | ***** | ***** | Iridium | nuclear marker |  | SBT/ Fluidigm | 201192A | 1:400 |
| 195Pt | ***** | ***** | ICSK1 | Cell Segmentation Kit 1 |  | SBT/ Fluidigm | Tis-00001 | 1:800 |
| 196Pt | ***** | ***** | ICSK2 | Cell Segmentation Kit 2 |  | SBT/ Fluidigm | Tis-00001 | 1:800 |
