## Supplementary material for "Spatial proteomic profiling of cardiac stromal populations and microenvironments in systemic sclerosis-associated primary myocardial involvement": Suppl.methods

**Supplementary Methods**

**Patient selection, biopsy assessment and clinical data**

Diagnostic biopsies from patients with SSc-VNM were obtained between January 2011 and June 2018 at three Italian SSc referral centres and University Hospital Erlangen. Biopsies from a-VNM and i-VNM were obtained between January 2016 and June 2018 at IRCCS San Raffaele Hospital, Milan. Clinical indications included dyspnoea, atypical chest pain with palpitations, syncope, lower-limb oedema, ventricular ectopy, non-sustained ventricular tachycardia or resuscitated cardiac arrest. Epidemiological, clinical and histopathological data are summarized in supplemental table S1. Biopsy indications were assessed in the context of contemporary endomyocardial-biopsy guidance.[45]

All biopsies were centrally reviewed by Prof. C. Basso, an expert cardiovascular pathologist at the University Hospital of Padua. Histological myocarditis was defined using the Dallas criteria. Viral causes were excluded by quantitative PCR for coxsackievirus, echovirus, adenovirus, influenza A/B, parvovirus B19, human herpesviruses 6-8, herpes simplex viruses 1/2, cytomegalovirus, varicella-zoster virus and Epstein-Barr virus. Histological fibrosis was quantified as the proportion of extracellular matrix in the assessed section stained with Masson’s Trichrome.

**Antibody validation and metal conjugation**

Unconjugated antibodies were evaluated by immunofluorescence on five µm paraffin sections of human myocardium using three serial dilutions. Antibodies showing the expected localization were conjugated to lanthanide or indium metals with the Maxpar X8 Antibody Labeling Kit (Standard BioTools) and retitrated by IMC. Commercially preconjugated antibodies were titrated directly by IMC. The complete antibody panel, isotope, clone, supplier, catalogue number and working dilution are provided in online supplemental table S2. The workflow followed established IMC antibody-validation and tissue-staining procedures.[46–48]

**Tissue staining**

Five-micrometre formalin-fixed, paraffin-embedded sections were incubated at 55°C for 60 min, deparaffinized in xylene (3×10 min), rehydrated through 100%, 90% and 80% ethanol (5 min each), rinsed in double-distilled water and washed in phosphate-buffered saline (PBS). Antigen retrieval was performed for 30 min at 96°C in Tris-EDTA-Tween buffer, pH 9.0. After approximately 20 min cooling at room temperature, sections were washed and blocked with 2% bovine serum albumin in PBS for 1 hour.

Sections were incubated overnight at 4°C with the antibody cocktail diluted in 2% bovine serum albumin/PBS. All study sections were stained with the same antibody master mix in a single batch. The next day, sections were washed in PBS containing 0.2% Tween-20 and then in PBS. Nuclear staining used iridium intercalator for 5 min. Sections were washed in PBS and double-distilled water, air-dried under low-dust conditions and stored at room temperature until acquisition.

**IMC acquisition and quality control**

Data were acquired on a Helios/Hyperion Imaging Mass Cytometer at the IMC Core Facility, University Hospital Erlangen, using CyTOF Software v7.0.8493. Standardized tuning followed each instrument restart, and calibration used a reference slide containing defined metal-isotope concentrations. Operating parameters were monitored against reference ranges. Regions of interest of approximately 1.5 mm² were ablated at 200 Hz, producing 1 µm² pixels. Raw data were retained as MCD and TXT files.

Two investigators independently reviewed each region in MCD Viewer v1.0560.6. Regions with insufficient staining or image quality were excluded. Marker-level heatmaps and multidimensional-scaling plots across samples were used to assess staining consistency and potential batch structure.

**Preprocessing, segmentation and signal normalization**

MCD/TXT files were converted to TIFF images with the readimc implementation in Steinbock v0.16.3, with hot-pixel removal. Multichannel cell segmentation used the Mesmer model in the DeepCell framework. Nuclear input combined Histone H3 and iridium DNA channels. Membrane/cytoplasmic input combined TNNT2, CD31, PDGFRα, PDGFRβ, CD45, CD68 and the two Cell Segmentation Kit channels. Segmentation masks and processed images were exported as OME-TIFF files, and per-cell intensities, morphology, centroid coordinates and image metadata were exported as tables.

Objects smaller than 30 or larger than 500 pixels were removed after visual validation of thresholds. Unnormalized per-cell mean intensities were transformed with the inverse hyperbolic sine function. Per-sample quantile normalization used the 0.95 reference quantile, followed by marker-level maximum or z-score rescaling as appropriate for the analysis. Data and spatial coordinates were assembled in a SpatialExperiment object with imcRtools.

**Cell phenotyping**

CD45^+^ immune and CD45^-^ non-immune cells were manually gated in 12 randomly selected regions with Cytomapper Shiny. These labels trained a random-forest classifier on quantile-normalized data that was applied to the remaining regions. Unsupervised clustering within each compartment used RPhenograph. Immune clustering used k=55 and TPSAB1, HLA-DR, CD34, CD14, CD163, CD20, CD86, CD4, CD8, CD3 and CD68. Non-immune clustering used k=40 and TNNT2, CD34, CD31, PDGFRα, α-SMA, PDGFRβ, ZEB1, POSTN, SOX9, PDPN, THY1, COL1, vWF, ADAM12, PI16, FSP1, TCF21 and FAP.

Cluster separation was visualized by UMAP. Annotation integrated scaled marker expression, established lineage-marker combinations and spatial localization in the source image. Cardiomyocyte clusters were merged into a single cardiomyocyte category. Clusters without a specific marker pattern were retained as undefined immune or undefined non-immune cells.

**Spatial analyses**

Pairwise cell-cell interactions were assessed with imcRtools on a four-nearest-neighbor graph. Observed neighbor counts were compared with 1000 label permutations. Positive significant interaction values denoted spatial enrichment and negative values spatial avoidance. For chord diagrams, reciprocal A-to-B and B-to-A scores were averaged within sample to yield a single undirected score and then summed across samples within disease group. These cohort-summed networks were used descriptively.

Cellular neighborhoods were generated by k-means clustering of local neighbor cell-type composition. Neighborhoods were named according to the dominant and distinguishing cell populations. For patient-level analyses, the frequency of a neighborhood was calculated as the percentage of all cells from that patient assigned to the neighborhood; patients with zero assigned cells were retained at 0%.

**Collagen quantification**

Collagen was segmented directly from the COL1 IMC channel. Per ROI, COL1 was arcsinh-transformed and Gaussian-smoothed (σ = 1 px; 1 px = 1 µm) and thresholded by Otsu; the median of the per-ROI Otsu thresholds was then applied as a single global threshold across all ROIs (with a 3-px morphological opening), yielding a binary collagen mask comparable across patients. For every segmented cell, two measures were read from this mask at the cell centroid: local collagen thickness (mean in-mask thickness within 20 µm) and distance to the nearest collagen-positive pixel. These were summarized to one value per patient and cell type (pooling cells across each patient's ROIs and requiring ≥ 5 cells) and compared across VNM groups at the patient level by Kruskal-Wallis tests with Holm-adjusted Dunn post-hoc tests. Figures show all cell types ranked by median (thickness, distance) or, for the between-group comparison, the cell types significant by Kruskal-Wallis or by any pairwise Dunn test, with significant pairs annotated.

**Statistical analysis**

All regions from a patient were pooled before calculating cell or neighborhood proportions, and patient was the inferential unit. Selected broad-compartment comparisons used Kruskal-Wallis tests followed by Dunn tests with Benjamini-Hochberg adjustment. Proportion analyses used the Propeller framework in speckle with arcsine-square-root-transformed proportions, robust empirical-Bayes variance estimation and no mean-variance trend fitting. Global FDR values were calculated across tested cell types or neighborhoods. Pairwise two-group Propeller analyses were adjusted across the three disease comparisons.

Fibrosis-group analyses primarily compared <10% versus ≥10% histological extracellular matrix, excluding patients without measurements. Continuous relationships with fibrosis used Spearman rank correlations. Correlation screens across multiple cell types were treated as exploratory and interpreted in relation to multiplicity correction. Statistical analyses were performed in R v4.3.2 using RStudio Desktop v2024.9.0.375.
