## Supplementary material for "Spatial proteomic profiling of cardiac stromal populations and microenvironments in systemic sclerosis-associated primary myocardial involvement": Suppl.Fig_legends

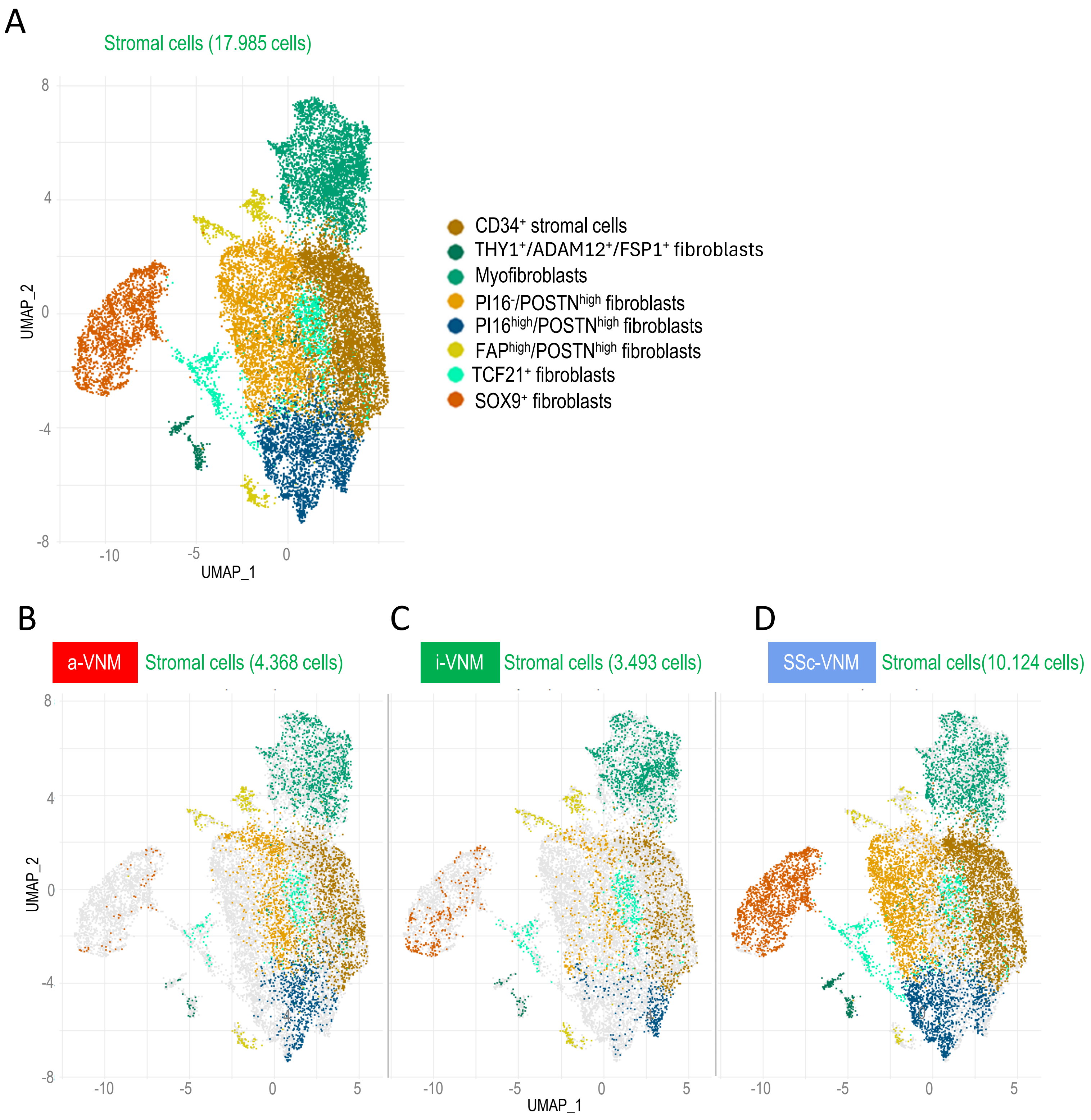


**Supplementary figure S1. IMC-resolved stromal-cell phenotypes.** (A) Uniform Manifold Approximation and Projection for Dimension Reduction (UMAP) illustrating all stromal cells across diseases. (B-D): UMAP illustrating the stromal cell clusters in a-VNM (B), i-VNM (C), and SSc-VNM (D). a-VNM, virus-negative myocarditis associated with a non-SSc systemic autoimmune rheumatic disease; i-VNM, isolated virus-negative myocarditis; SSc-VNM, systemic sclerosis-associated virus-negative myocarditis.

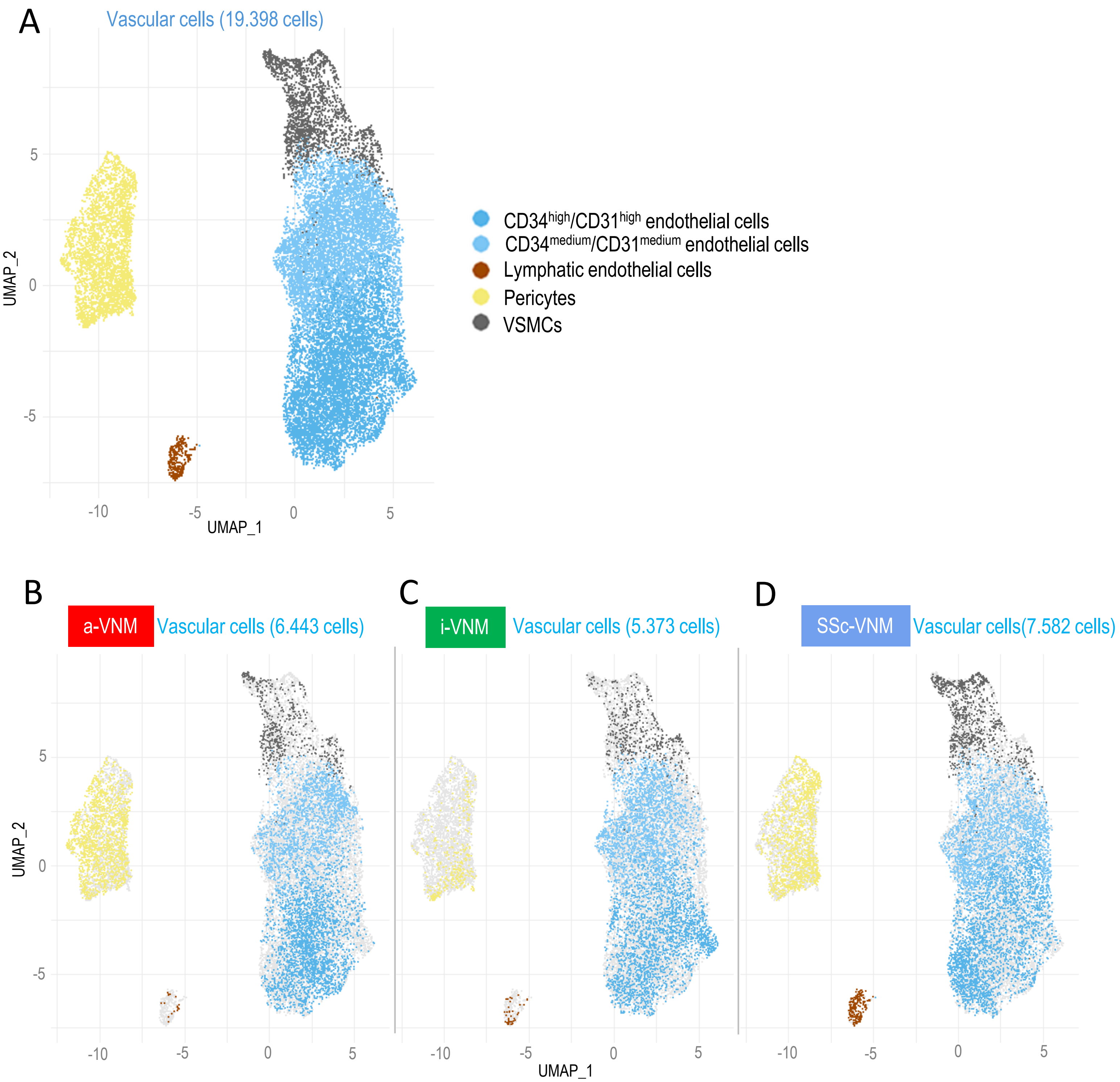


**Supplementary figure S2. IMC-resolved vascular-cell phenotypes.** (A) Uniform Manifold Approximation and Projection for Dimension Reduction (UMAP) illustrating all vascular cells across diseases. (B-D): UMAP illustrating the vascular cell clusters in a-VNM (B), i-VNM (C), and SSc-VNM (D). a-VNM, virus-negative myocarditis associated with a non-SSc systemic autoimmune rheumatic disease; i-VNM, isolated virus-negative myocarditis; SSc-VNM, systemic sclerosis-associated virus-negative myocarditis, VSMCs, vascular smooth muscle cells.

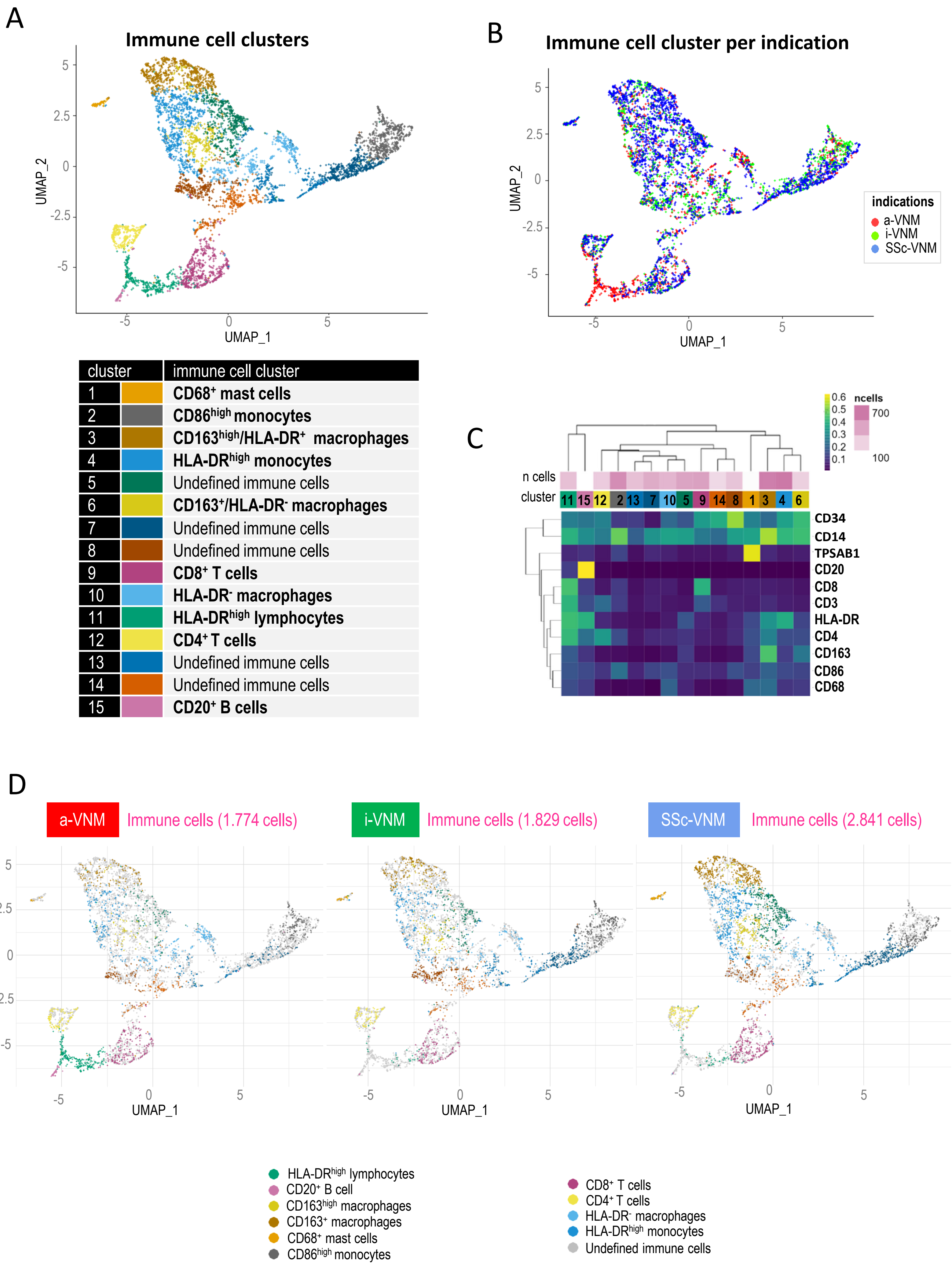
A
Immune cell clusters
UMAP_2
UMAP_1
cluster
immune cell cluster
1 CD68+ mast cells
2 CD86high monocytes
3 CD163high/HLA-DR+ macrophages
4 HLA-DRhigh monocytes
5 Undefined immune cells
6 CD163+/HLA-DR- macrophages
7 Undefined immune cells
8 Undefined immune cells
9 CD8+ T cells
10 HLA-DR- macrophages
11 HLA-DRhigh lymphocytes
12 CD4+ T cells
13 Undefined immune cells
14 Undefined immune cells
15 CD20+ B cells
B
Immune cell cluster per indication
indications
a-VNM
i-VNM
SSc-VNM
C
n cells
cluster
11 15 12 2 13 7 10 5 9 14 8 1 3 4 6
ncells
700
100
CD34
CD14
TPSAB1
CD20
CD8
CD3
HLA-DR
CD4
CD163
CD86
CD68
D
a-VNM
Immune cells (1.774 cells)
i-VNM
Immune cells (1.829 cells)
SSc-VNM
Immune cells (2.841 cells)
HLA-DRhigh lymphocytes
CD20+ B cell
CD163high macrophages
CD163+ macrophages
CD68+ mast cells
CD86high monocytes
CD8+ T cells
CD4+ T cells
HLA-DR- macrophages
HLA-DRhigh monocytes
Undefined immune cells

**Supplementary figure S3. Phenotypic characterization of CD45$^+$ immune populations.** (A) Uniform Manifold Approximation and Projection for Dimension Reduction (UMAP) of immune clusters 1-15. (B) UMAP coloured by disease. (C) Heatmap of marker expression across clusters. (D) Phenotype-coloured UMAPs separated by a-VNM, i-VNM and SSc-VNM. a-VNM, virus-negative myocarditis associated with a non-SSc systemic autoimmune rheumatic disease; i-VNM, isolated virus-negative myocarditis; SSc-VNM, systemic sclerosis-associated virus-negative myocarditis.

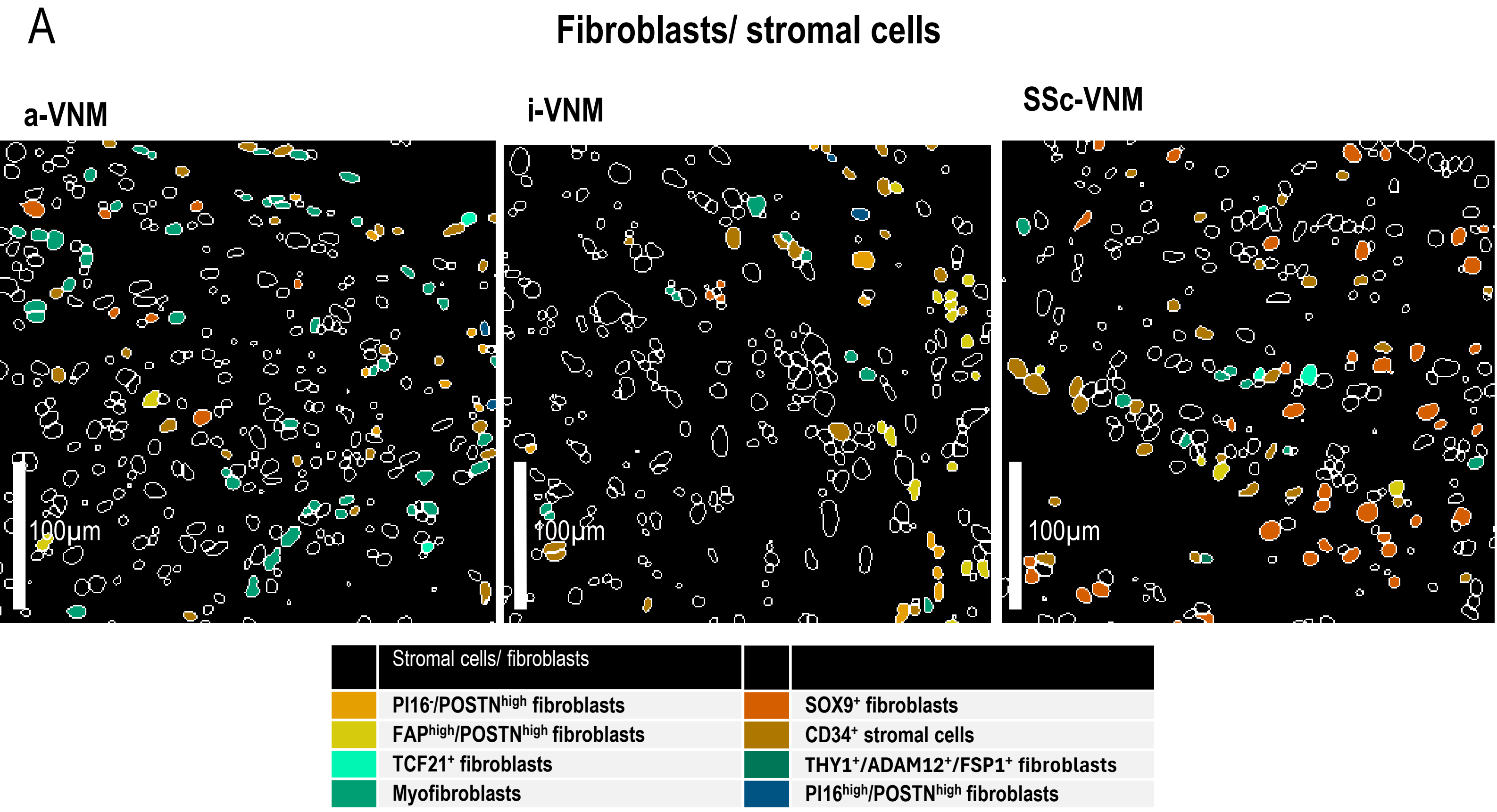
A
Fibroblasts/ stromal cells
a-VNM
i-VNM
SSc-VNM
100µm
100µm
100µm
Stromal cells/ fibroblasts
PI16⁻/POSTNʰⁱᵍʰ fibroblasts
FAPʰⁱᵍʰ/POSTNʰⁱᵍʰ fibroblasts
TCF21⁺ fibroblasts
Myofibroblasts
SOX9⁺ fibroblasts
CD34⁺ stromal cells
THY1⁺/ADAM12⁺/FSP1⁺ fibroblasts
PI16ʰⁱᵍʰ/POSTNʰⁱᵍʰ fibroblasts

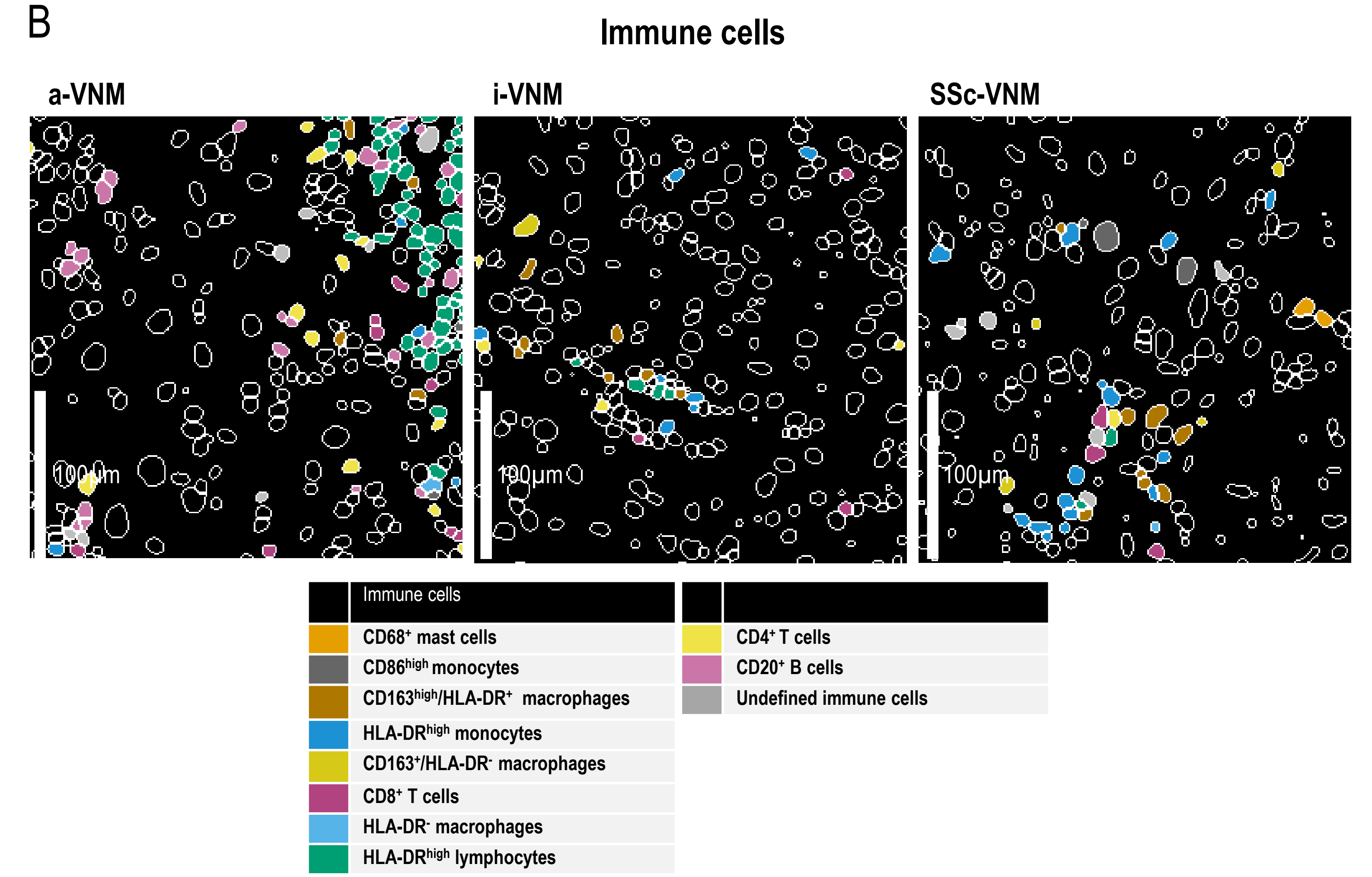
B
Immune cells
a-VNM
i-VNM
SSc-VNM
100µm
100µm
100µm
Immune cells
CD68⁺ mast cells
CD86ʰⁱᵍʰ monocytes
CD163ʰⁱᵍʰ/HLA-DR⁺ macrophages
HLA-DRʰⁱᵍʰ monocytes
CD163⁺/HLA-DR⁻ macrophages
CD8⁺ T cells
HLA-DR⁻ macrophages
HLA-DRʰⁱᵍʰ lymphocytes
CD4⁺ T cells
CD20⁺ B cells
Undefined immune cells

**Supplementary figure S4. Representative cytomapper images of fibroblast/stromal cell (A) and immune cell (B) subsets spatial distribution across myocarditis entities.** Bars represent 100 µm. a-VNM, virus-negative myocarditis associated with a non-SSc systemic autoimmune rheumatic disease; i-VNM, isolated virus-negative myocarditis; SSc-VNM, systemic sclerosis-associated virus-negative myocarditis.

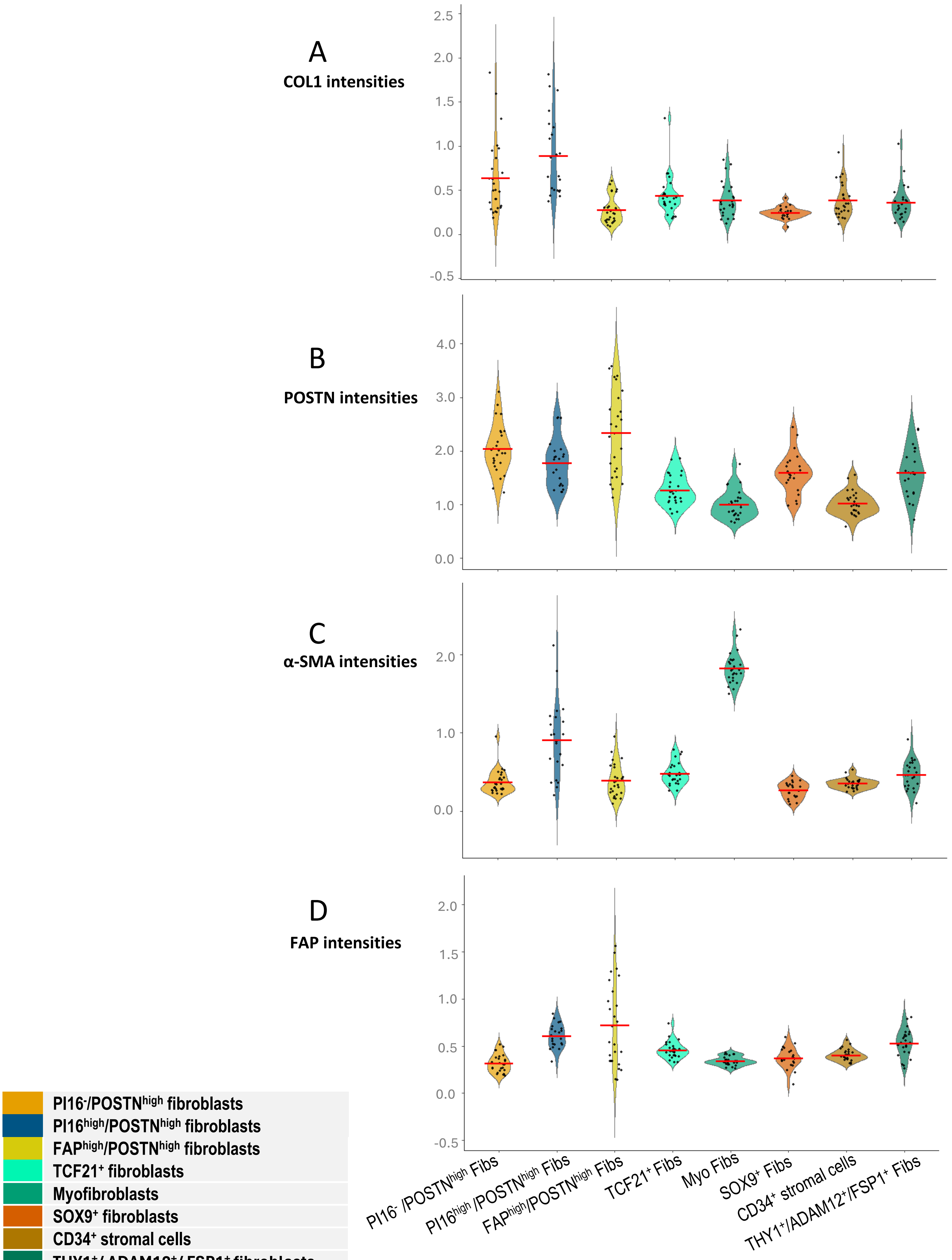
A
COL1 intensities
B
POSTN intensities
C
α-SMA intensities
D
FAP intensities
PI16-/POSTNhigh fibroblasts
PI16high/POSTNhigh fibroblasts
FAPhigh/POSTNhigh fibroblasts
TCF21+ fibroblasts
Myofibroblasts
SOX9+ fibroblasts
CD34+ stromal cells
THY1+/ ADAM12+/ FSP1+ fibroblasts
PI16- /POSTNhigh Fibs
PI16high /POSTNhigh Fibs
FAPhigh/POSTNhigh Fibs
TCF21+ Fibs
Myo Fibs
SOX9+ Fibs
CD34+ stromal cells
THY1+/ADAM12+/FSP1+ Fibs

**Supplementary figure S5. Fibroblast-marker expression across stromal cell phenotypes.** Violin plots illustrating patient-level mean arcsinh-transformed IMC intensities for (A) COL1A1, (B) POSTN, (C) α-smooth muscle actin and (D) FAP across stromal phenotypes. Points are patients, violins show the distribution and red bars indicate group means.

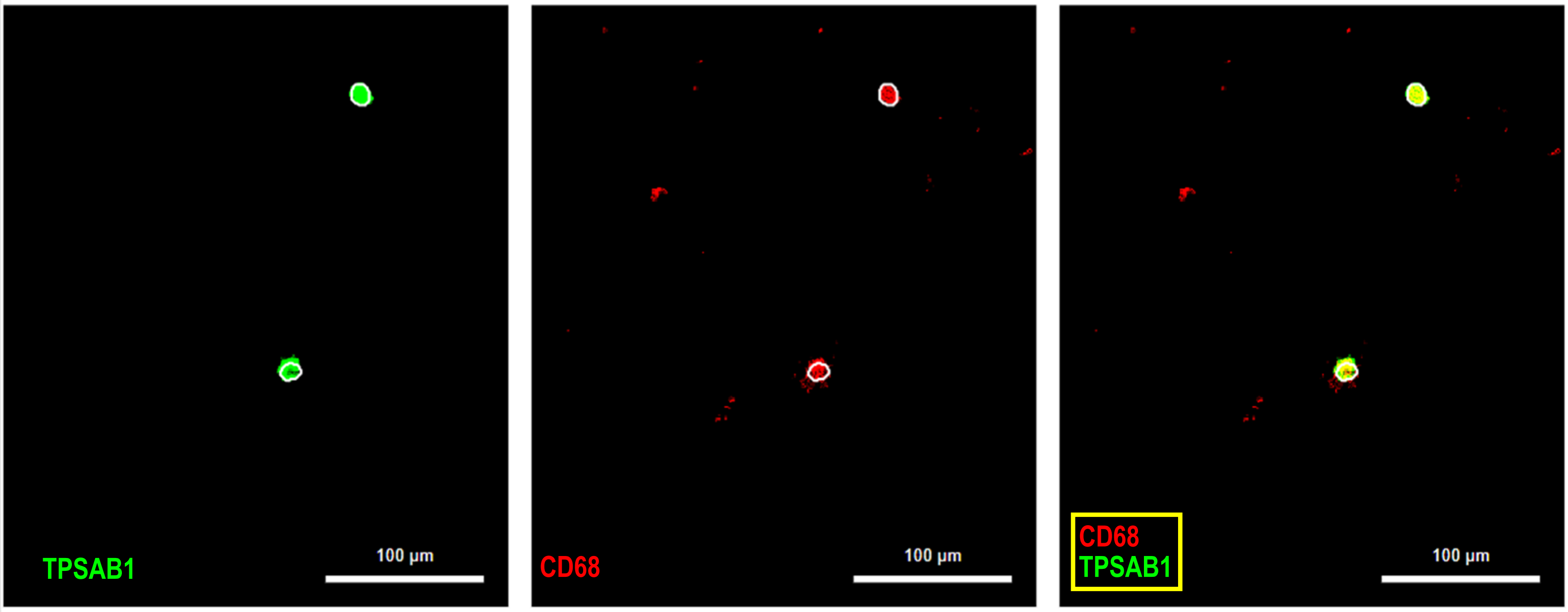


**Supplementary figure S6. TPSAB1⁺/CD68⁺ mast cells.** Representative multiplexed image of TPSAB1 (green), CD68 (red) and merged image.  Scale bars are 100 µm.

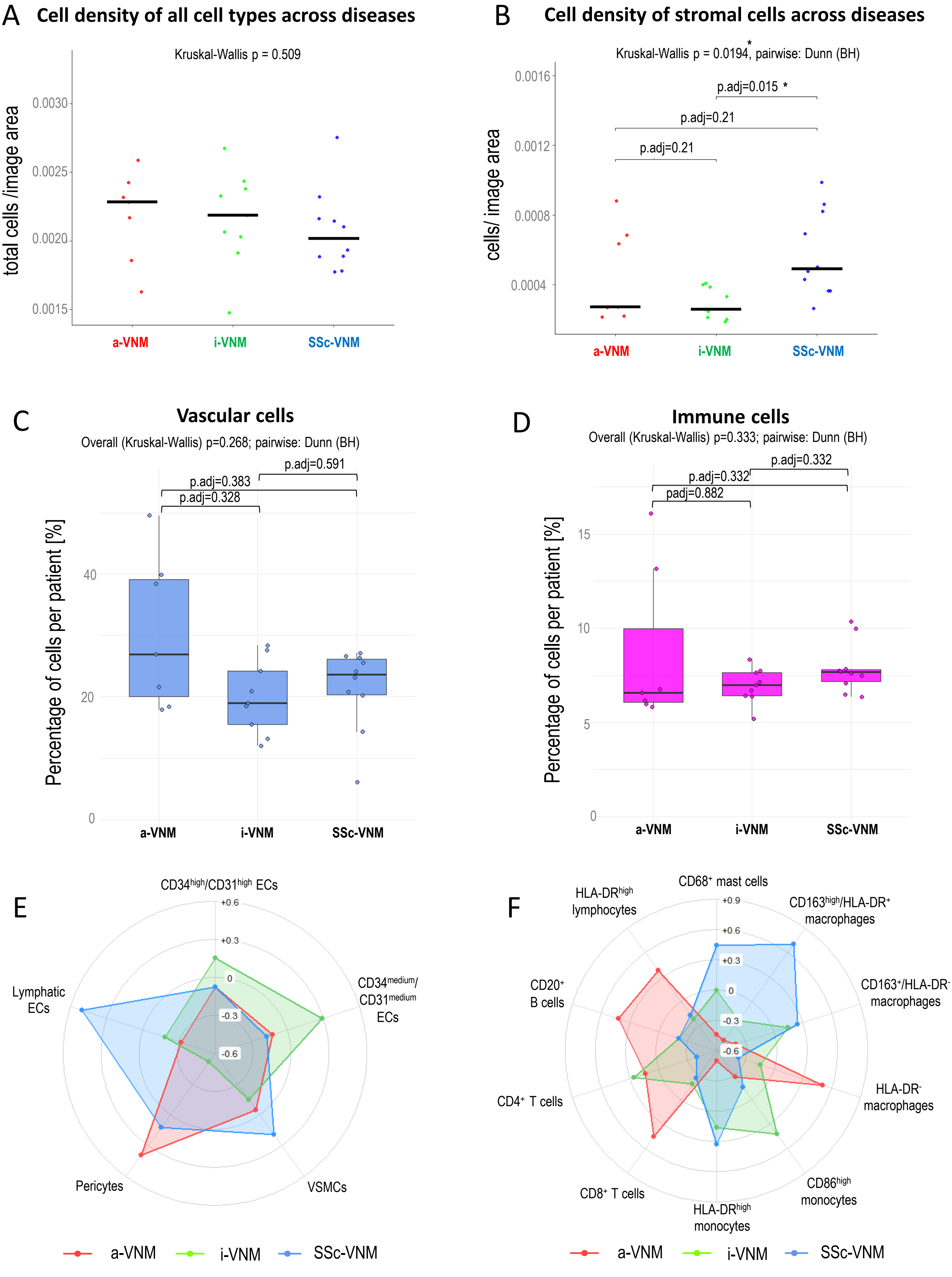

A
Cell density of all cell types across diseases
Kruskal-Wallis p = 0.509
total cells /image area
0.0030
0.0025
0.0020
0.0015
a-VNM
i-VNM
SSc-VNM
B
Cell density of stromal cells across diseases
Kruskal-Wallis p = 0.0194*, pairwise: Dunn (BH)
p.adj=0.015 *
p.adj=0.21
p.adj=0.21
cells/ image area
0.0016
0.0012
0.0008
0.0004
a-VNM
i-VNM
SSc-VNM
C
Vascular cells
Overall (Kruskal-Wallis) p=0.268; pairwise: Dunn (BH)
p.adj=0.591
p.adj=0.383
p.adj=0.328
Percentage of cells per patient [%]
40
20
0
a-VNM
i-VNM
SSc-VNM
D
Immune cells
Overall (Kruskal-Wallis) p=0.333; pairwise: Dunn (BH)
p.adj=0.332
p.adj=0.332
padj=0.882
Percentage of cells per patient [%]
15
10
5
0
a-VNM
i-VNM
SSc-VNM
E
CD34high/CD31high ECs
+0.6
+0.3
0
-0.3
-0.6
CD34medium/ CD31medium ECs
VSMCs
Pericytes
Lymphatic ECs
a-VNM
i-VNM
SSc-VNM
F
CD68+ mast cells
+0.9
+0.6
+0.3
0
-0.3
-0.6
CD163high/HLA-DR+ macrophages
CD163+/HLA-DR- macrophages
HLA-DR- macrophages
CD86high monocytes
HLA-DRhigh monocytes
CD8+ T cells
CD4+ T cells
CD20+ B cells
HLA-DRhigh lymphocytes
a-VNM
i-VNM
SSc-VNM

**Supplementary figure S7. Cell density and broad compartment proportions across diseases.** (A) Total-cell density across myocarditis entities. (B) Stromal-cell density across disease entities. (C-D) Patient-level proportions of (C vascular cells and (D) immune cells. Points are patients; boxplots show medians and interquartile ranges. Z-scaled radar charts showing the distribution of (E) vascular cells and (F) immune cells across myocarditis entities. a-VNM, virus-negative myocarditis associated with a non-SSc systemic autoimmune rheumatic disease; i-VNM, isolated virus-negative myocarditis; SSc-VNM, systemic sclerosis-associated virus-negative myocarditis.

.

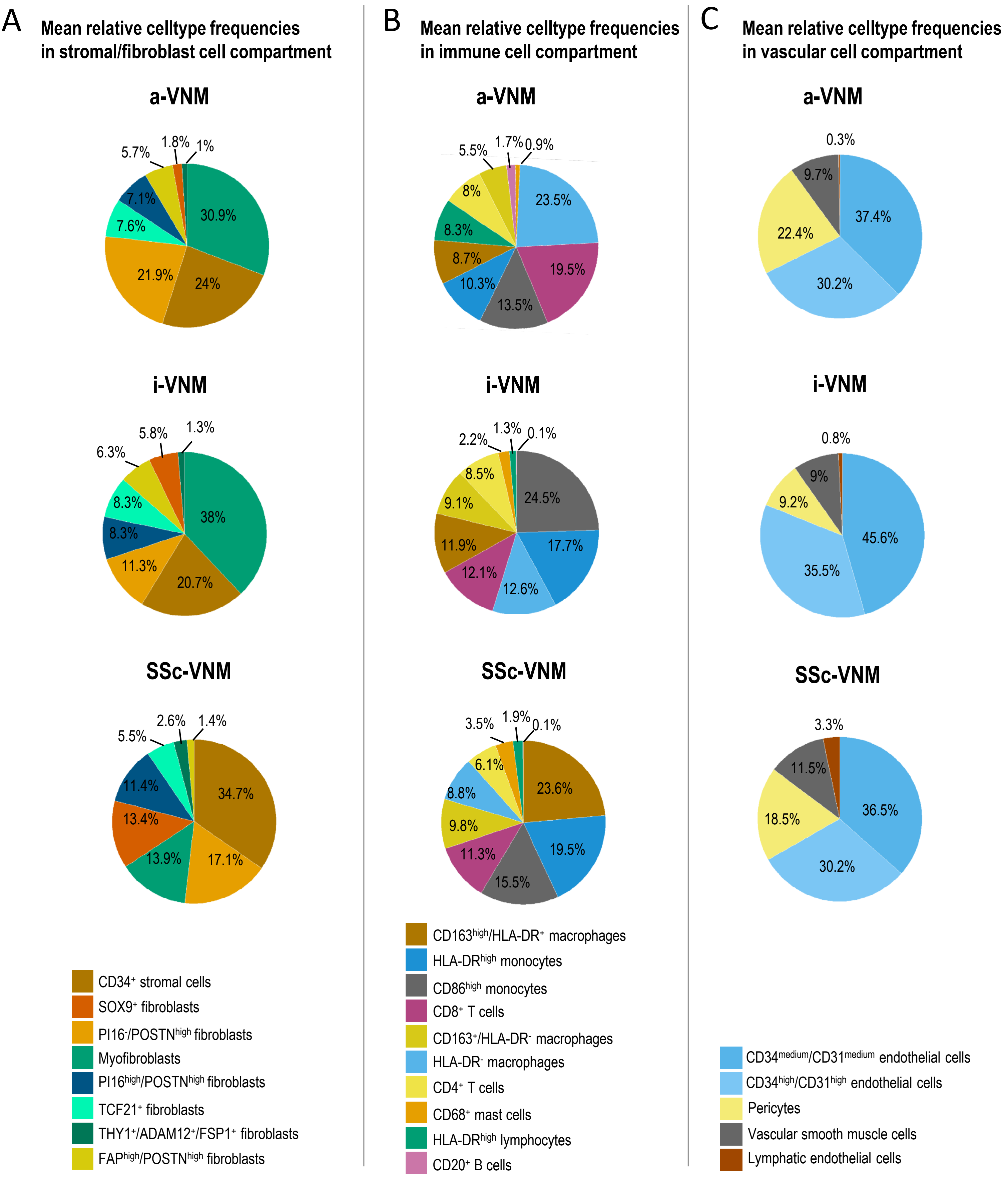


**Supplementary figure S8. Compartment-specific cellular composition across diseases.** Pie charts illustrating the mean phenotype proportions within the (A) stromal, (B) immune and (C) vascular compartments for SSc-VNM, a-VNM and i-VNM. a-VNM, virus-negative myocarditis associated with a non-SSc systemic autoimmune rheumatic disease; i-VNM, isolated virus-negative myocarditis; SSc-VNM, systemic sclerosis-associated virus-negative myocarditis.

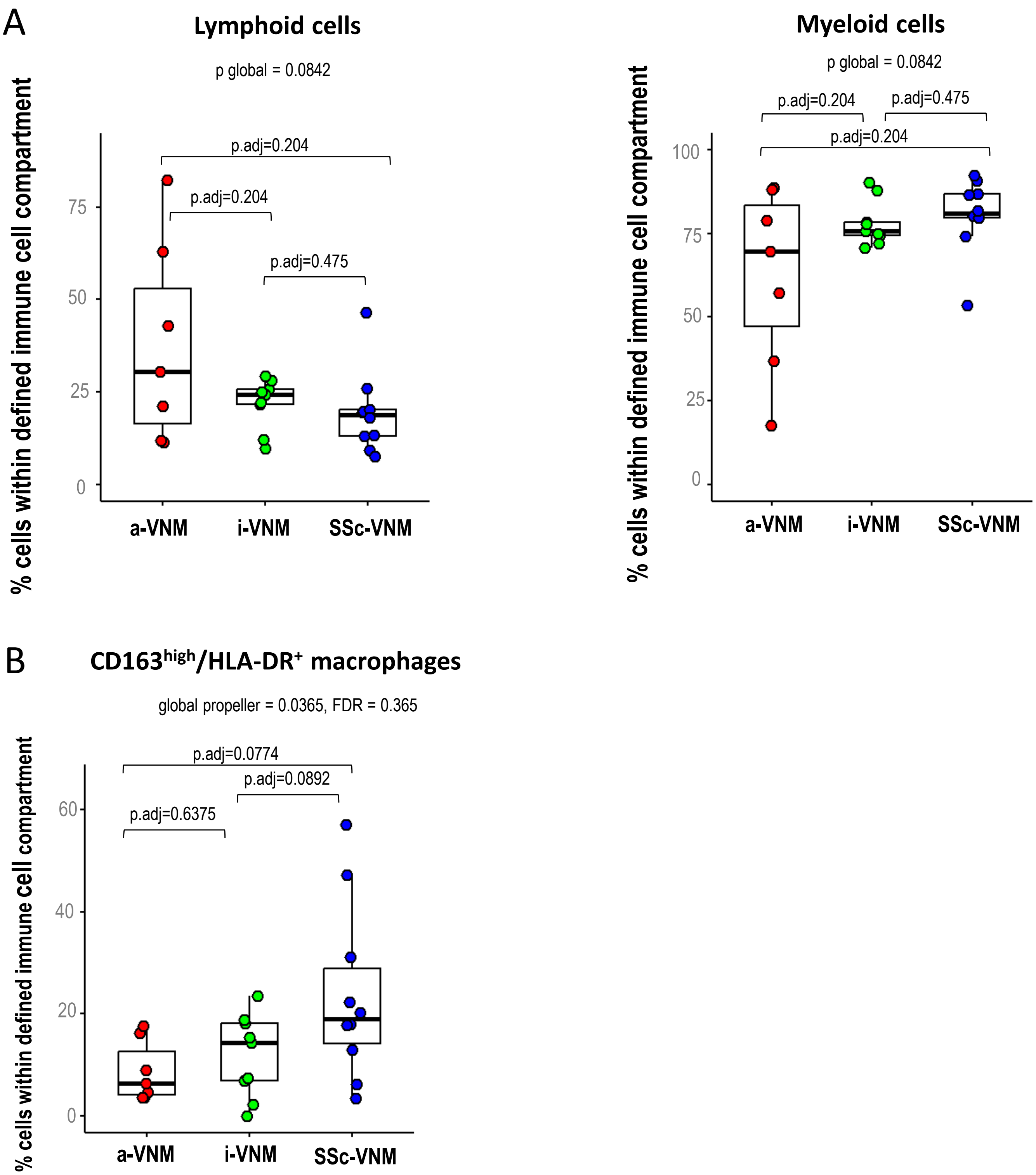


**Supplementary figure S9. Immune-cell proportions across diseases.** (A) Lymphoid and myeloid proportions among defined immune cells. (B) CD163$^{high}$/HLA-DR$^{+}$ macrophages. Boxplots show patient-level distributions. a-VNM, virus-negative myocarditis associated with a non-SSc systemic autoimmune rheumatic disease; i-VNM, isolated virus-negative myocarditis; SSc-VNM, systemic sclerosis-associated virus-negative myocarditis.

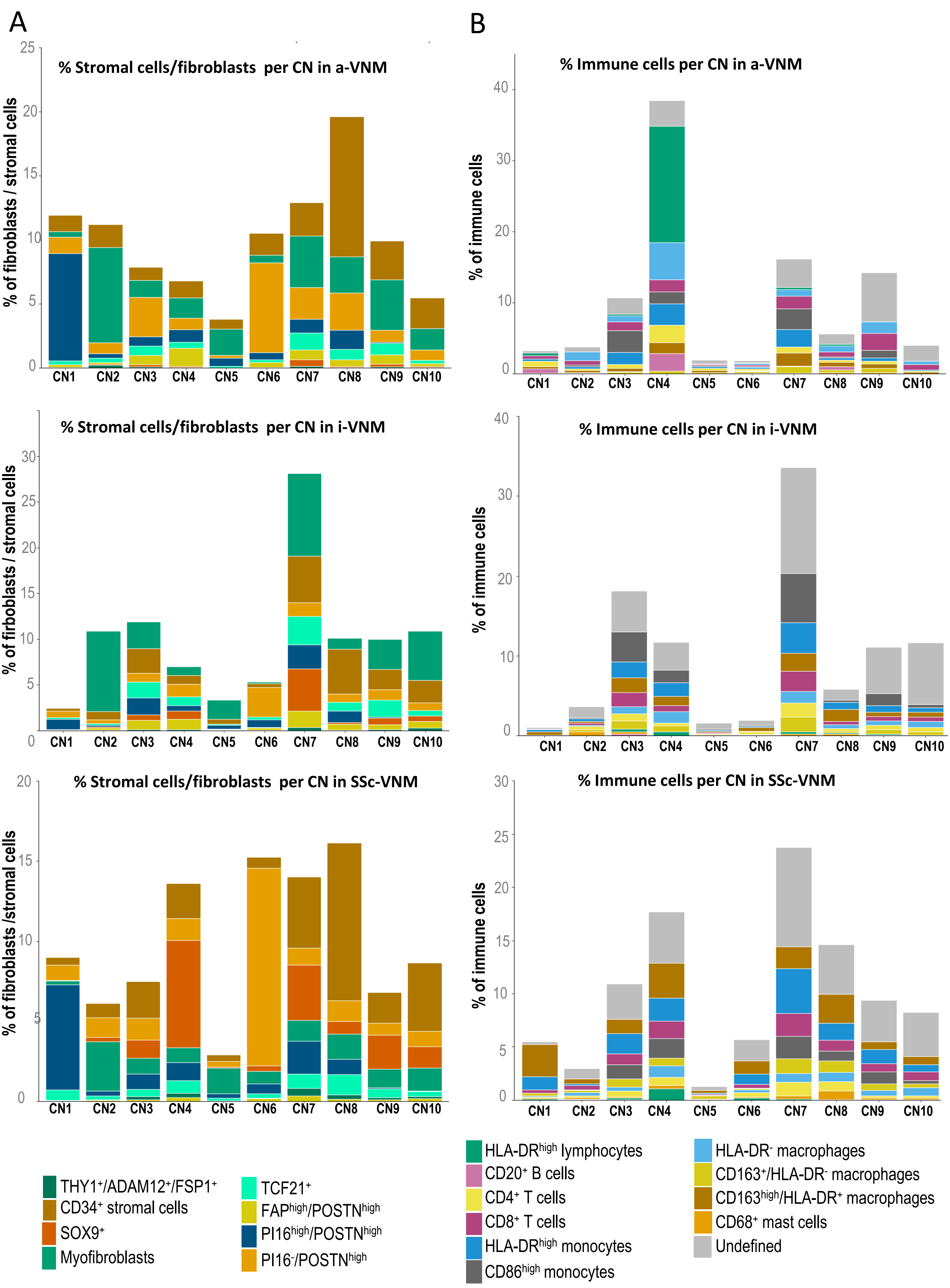

A
% Stromal cells/fibroblasts per CN in a-VNM
% of fibroblasts / stromal cells
% Stromal cells/fibroblasts per CN in i-VNM
% of firboblasts / stromal cells
% Stromal cells/fibroblasts per CN in SSc-VNM
% of fibroblasts /stromal cells
CN1 CN2 CN3 CN4 CN5 CN6 CN7 CN8 CN9 CN10
B
% Immune cells per CN in a-VNM
% Immune cells per CN in i-VNM
% Immune cells per CN in SSc-VNM
% of immune cells
THY1+/ADAM12+/FSP1+
CD34+ stromal cells
SOX9+
Myofibroblasts
TCF21+
FAPhigh/POSTNhigh
PI16high/POSTNhigh
PI16-/POSTNhigh
HLA-DRhigh lymphocytes
CD20+ B cells
CD4+ T cells
CD8+ T cells
HLA-DRhigh monocytes
CD86high monocytes
HLA-DR- macrophages
CD163+/HLA-DR- macrophages
CD163high/HLA-DR+ macrophages
CD68+ mast cells
Undefined

**Supplementary figure S10. Cell-type composition of cellular neighborhoods.** Stacked bars show the relative proportion of celltypes within the (A) fibroblast/stromal and (B) immune cell compartment across cellular neighborhood (CN) and diseases.

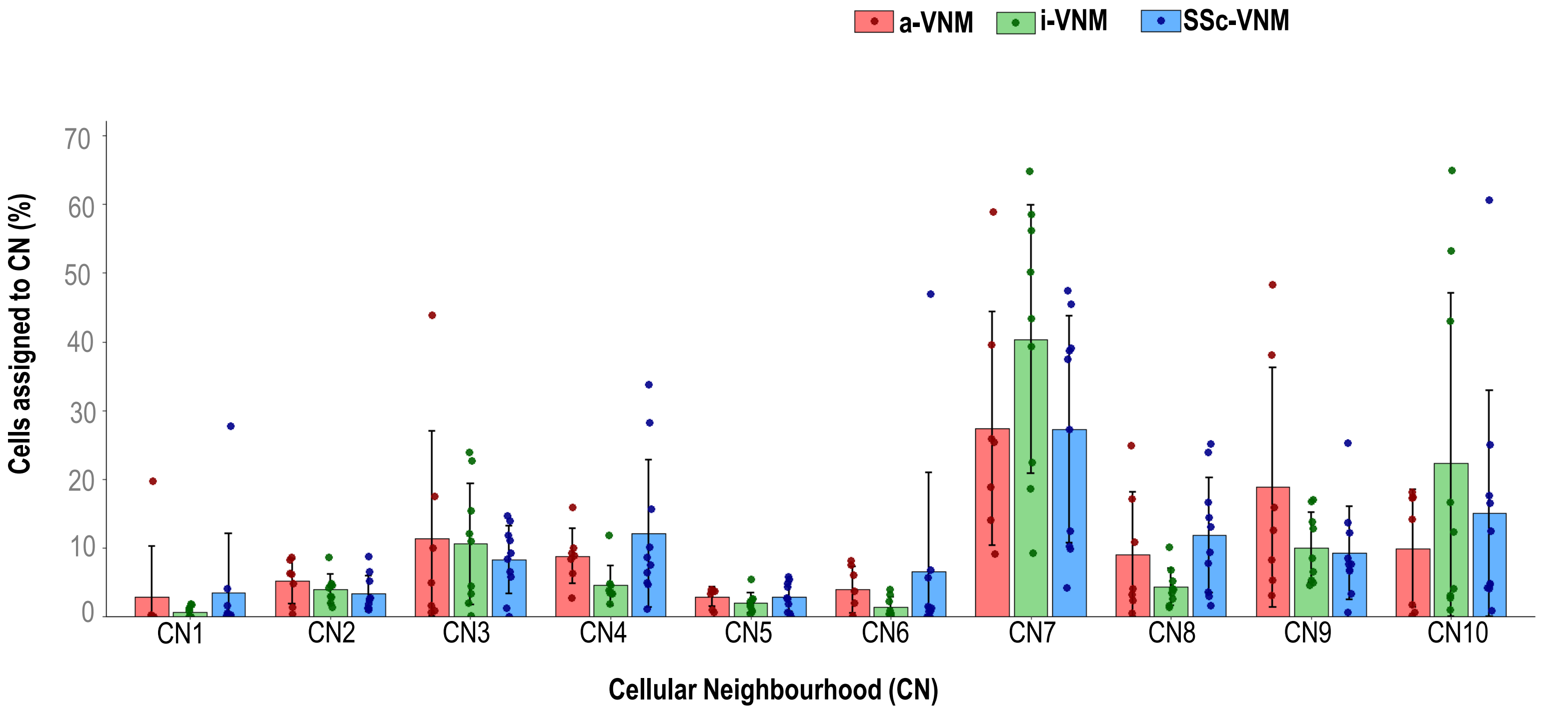


**Supplementary figure S11. Total cell distribution across cellular neighborhoods by disease.** Patient-level cell-type frequencies assigned to each CN are shown for a-VNM, i-VNM and SSc-VNM. a-VNM, virus-negative myocarditis associated with a non-SSc systemic autoimmune rheumatic disease; i-VNM, isolated virus-negative myocarditis; SSc-VNM, systemic sclerosis-associated virus-negative myocarditis.

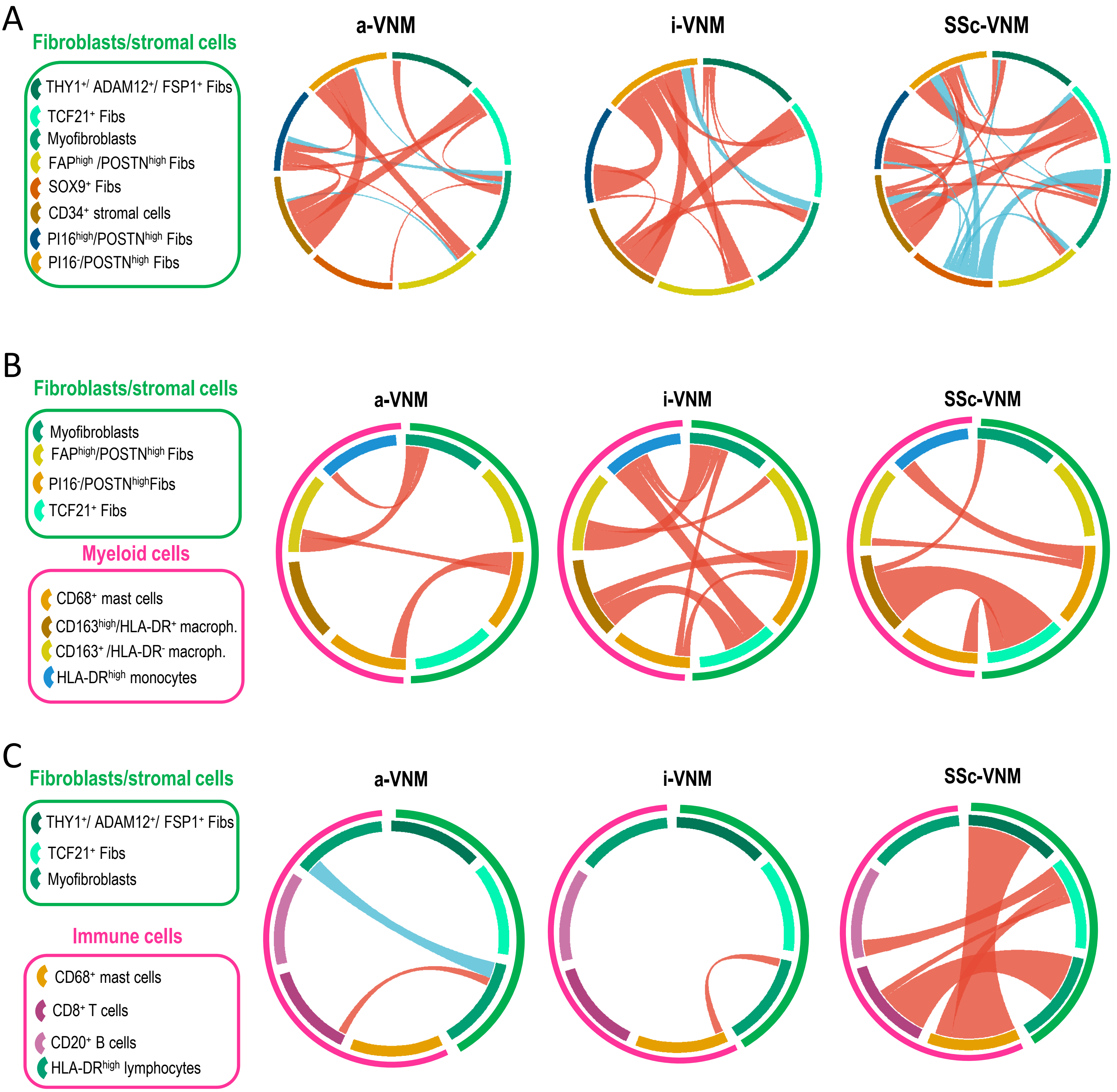
A
Fibroblasts/stromal cells
THY1+/ ADAM12+/ FSP1+ Fibs
TCF21+ Fibs
Myofibroblasts
FAPhigh /POSTNhigh Fibs
SOX9+ Fibs
CD34+ stromal cells
PI16high/POSTNhigh Fibs
PI16-/POSTNhigh Fibs
a-VNM
i-VNM
SSc-VNM
B
Fibroblasts/stromal cells
Myofibroblasts
FAPhigh/POSTNhigh Fibs
PI16-/POSTNhighFibs
TCF21+ Fibs
Myeloid cells
CD68+ mast cells
CD163high/HLA-DR+ macroph.
CD163+ /HLA-DR- macroph.
HLA-DRhigh monocytes
a-VNM
i-VNM
SSc-VNM
C
Fibroblasts/stromal cells
THY1+/ ADAM12+/ FSP1+ Fibs
TCF21+ Fibs
Myofibroblasts
Immune cells
CD68+ mast cells
CD8+ T cells
CD20+ B cells
HLA-DRhigh lymphocytes
a-VNM
i-VNM
SSc-VNM

**Supplementary figure S12. Spatial cell-cell interaction networks across myocarditis entities.** Chord diagrams showing significant permutation-based interactions between fibroblasts/stromal cells with fibroblasts/stromal cells (A), myeloid cells (B), or immune cells (C) a-VNM, i-VNM and SSc-VNM using four-nearest-neighbor graphs. Bidirectional interaction scores were averaged per sample and summed within disease group. Red denotes spatial enrichment and blue spatial avoidance; chord width is proportional to the absolute summed interaction score. a-VNM, virus-negative myocarditis associated with a non-SSc systemic autoimmune rheumatic disease; i-VNM, isolated virus-negative myocarditis; SSc-VNM, systemic sclerosis-associated virus-negative myocarditis.

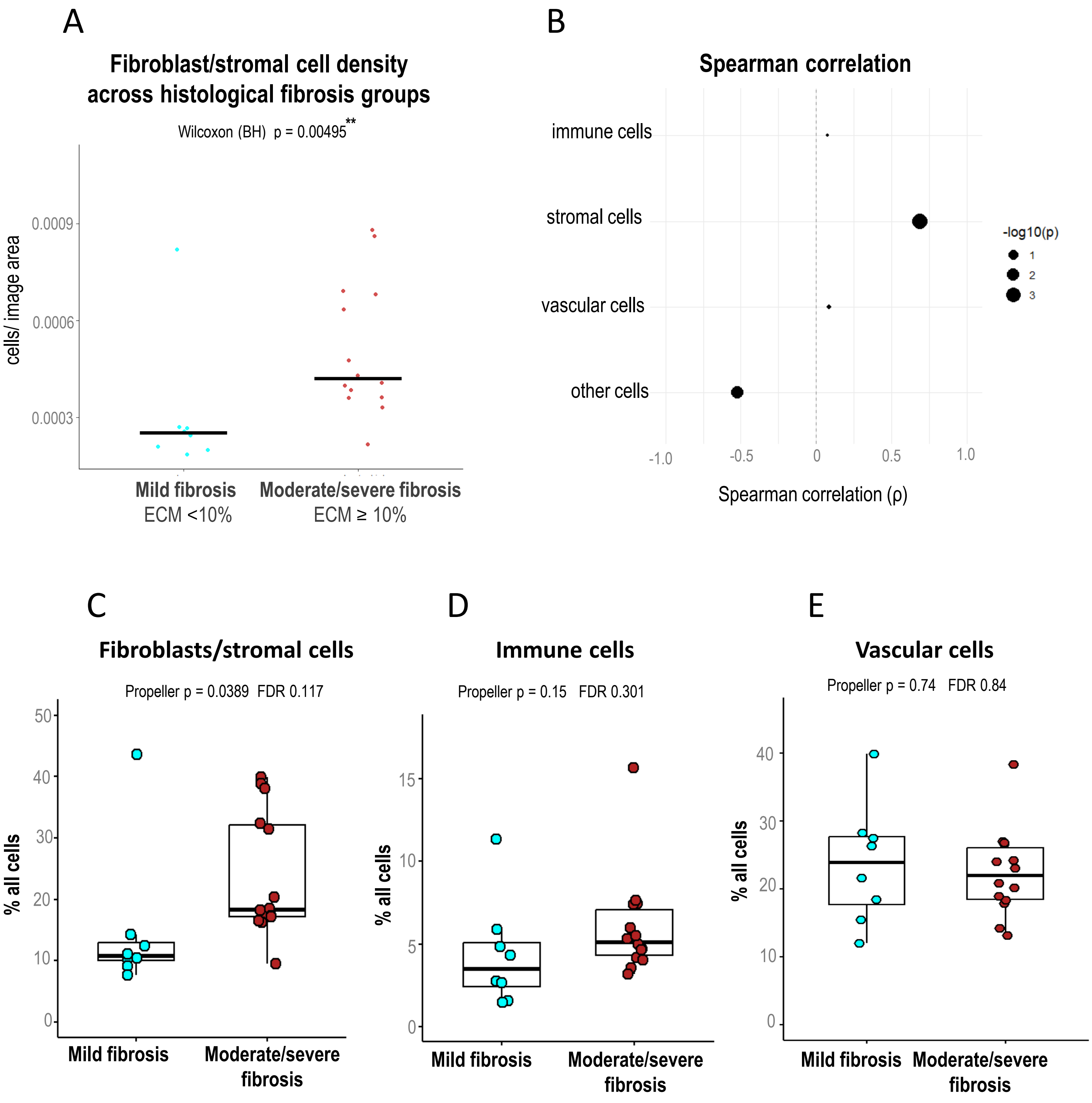


**Supplementary figure S13. Broad cellular compartments across myocarditis with mild or moderate/severe histological fibrosis.** (A) Mean patient-level fibroblast/stromal cell density in mild and moderate/severe fibrosis. (B) Spearman correlations between continuous ECM percentage and broad cell-compartment densities. (C-E) Boxplots showing the stromal cell (C), defined immune-cell (D), and vascular cell (E) proportions across fibrosis groups.

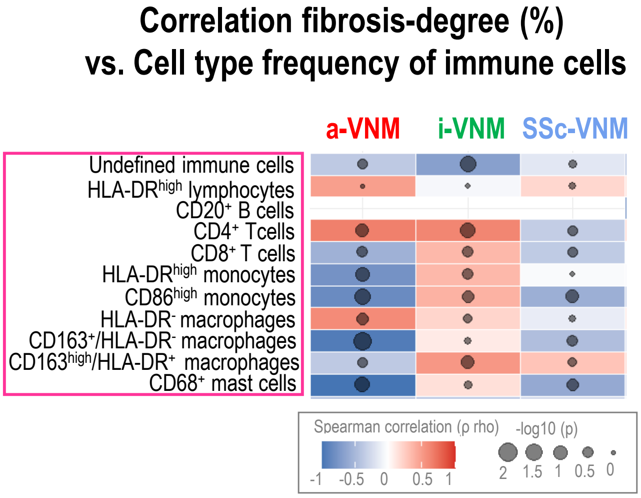


**Supplementary figure S14. Exploratory correlations between histological fibrosis and immune-cell frequencies.** Heatmap illustrating the disease-specific Spearman rho and p values for a-VNM (n=6), i-VNM (n=9) and SSc-VNM (n=7). a-VNM, virus-negative myocarditis associated with a non-SSc systemic autoimmune rheumatic disease; i-VNM, isolated virus-negative myocarditis; SSc-VNM, systemic sclerosis-associated virus-negative myocarditis.

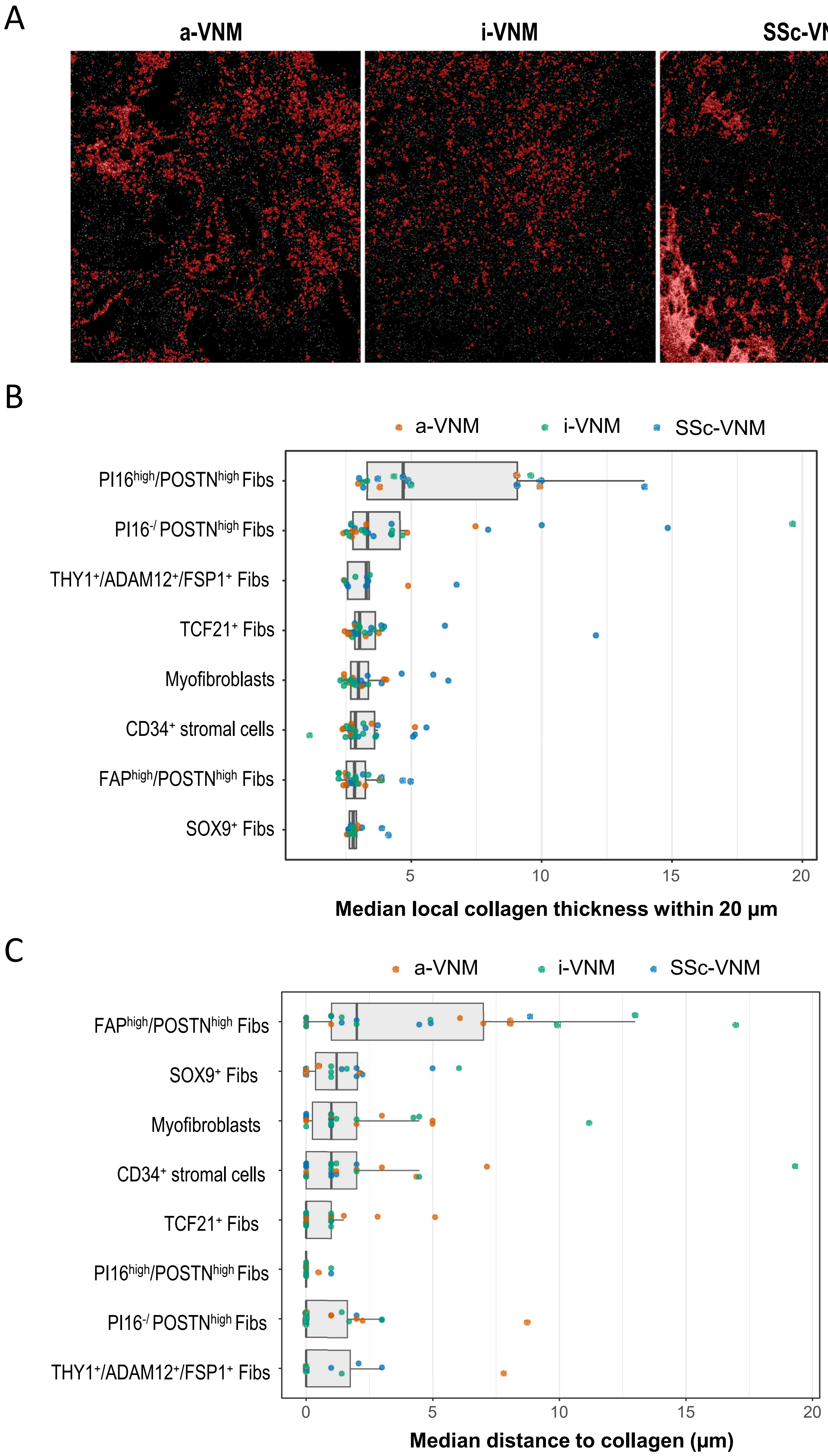
A
a-VNM
i-VNM
SSc-VNM
B
a-VNM
i-VNM
SSc-VNM
PI16high/POSTNhigh Fibs
PI16-/ POSTNhigh Fibs
THY1+/ADAM12+/FSP1+ Fibs
TCF21+ Fibs
Myofibroblasts
CD34+ stromal cells
FAPhigh/POSTNhigh Fibs
SOX9+ Fibs
5
10
15
20
Median local collagen thickness within 20 μm
C
a-VNM
i-VNM
SSc-VNM
FAPhigh/POSTNhigh Fibs
SOX9+ Fibs
Myofibroblasts
CD34+ stromal cells
TCF21+ Fibs
PI16high/POSTNhigh Fibs
PI16-/ POSTNhigh Fibs
THY1+/ADAM12+/FSP1+ Fibs
0
5
10
15
20
Median distance to collagen (μm)

**Supplementary figure S15. Descriptive overview of patient-level fibroblast/stromal cell-type specific collagen thickness and distance to collagen signal.** (A) Representative collagen masks. (B) Local collagen thickness. Median collagen thickness within 20 µm of each cell type (local diameter of nearby collagen deposits). (C) Distance to nearest collagen signal. Median distance (µm) from each cell type to the nearest collagen-positive pixel. Cell types are coloured by compartment. Descriptive overviews (no tests). Only patients with ≥ 5 cells of that specific cell type are shown.

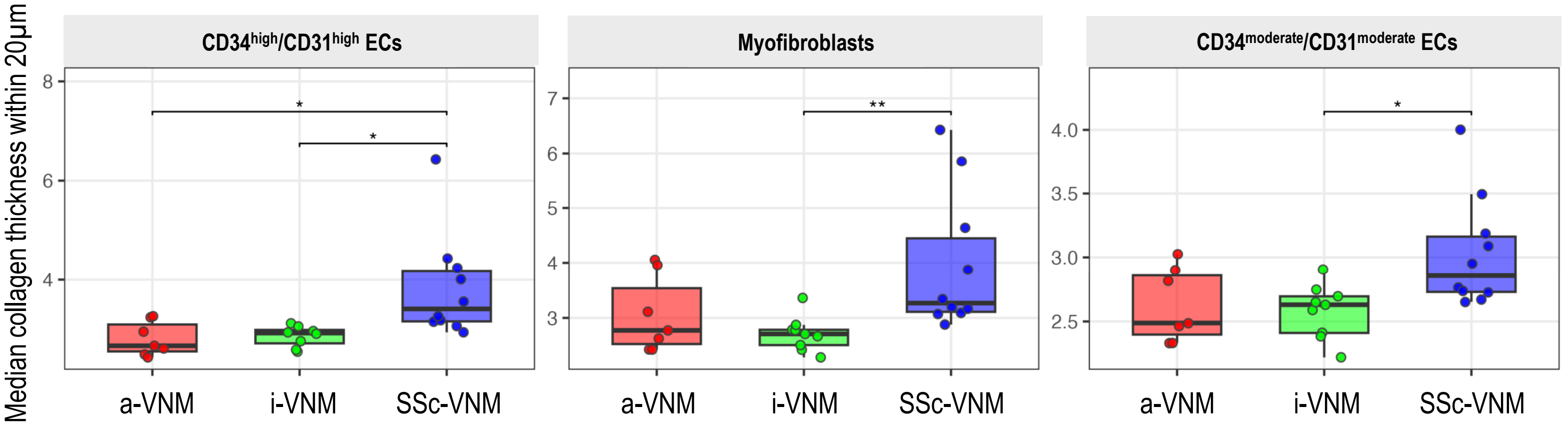


**Supplementary figure S16. Patient-level cell-type specific median within 20 µm collagen thickness across myocarditis entities.** Kruskal-Wallis tests with Holm-adjusted Dunn post-hoc tests were applied. Brackets mark significant Dunn pairs. Each point is one patient. Only patients with ≥ 5 cells of that specific cell type are shown. * p< 0.05; ** p<0.01. a-VNM, virus-negative myocarditis associated with a non-SSc systemic autoimmune rheumatic disease; i-VNM, isolated virus-negative myocarditis; SSc-VNM, systemic sclerosis-associated virus-negative myocarditis.
